# Developmentally regulated PERK activity renders dendritic cells insensitive to subtilase cytotoxin-induced integrated stress response

**DOI:** 10.1101/2020.04.27.063438

**Authors:** Andreia Mendes, Julien P. Gigan, Christian Rodriguez Rodrigues, Sébastien A. Choteau, Doriane Sanseau, Daniela Barros, Catarina Almeida, Voahirana Camosseto, Rafael J. Argüello, Lionel Chasson, Adrienne W. Paton, James C. Paton, Ana-Maria Lennon-Duménil, Evelina Gatti, Philippe Pierre

## Abstract

In stressed cells, phosphorylation of eukaryotic initiation factor 2α (eIF2α) controls transcriptome-wide changes in mRNA translation and gene expression known as the integrated stress response (ISR). We show here that dendritic cells (DCs) display unusually high eIF2α phosphorylation, which is mostly caused by a developmentally regulated activation of the ER kinase PERK (EIF2AK3). Despite high p-eIF2α levels, differentiated DCs display active protein synthesis and no signs of a chronic ISR. eIF2α phosphorylation does not majorly impact DC differentiation nor cytokines production. It is however important to adapt protein homeostasis to the variations imposed on DCs by the immune or physiological contexts. This biochemical specificity prevents translation arrest and expression of the transcription factor ATF4 during ER-stress induction by subtilase cytotoxin or upon DC stimulation with bacterial lipopolysaccharides. This is also exemplified by the influence of the actin cytoskeleton dynamics on eIF2α phosphorylation and the migratory deficit observed in PERK-deficient DCs.

## Introduction

Dendritic cells (DC) are key regulators of both protective immune responses and tolerance to self-antigens (Dalod et al., 2014). DCs are professional antigen presenting cells (APC), equipped with pattern recognition receptors (PRRs), capable of recognizing microbe-associated molecular patterns (MAMPs) (Akira et al., 2006) and enhance their immuno-stimulatory activity (Steinman, 2007). MAMPs detection by DCs triggers the process of maturation/activation, that culminates in the unique capacity of priming naïve T cells in secondary lymphoid organs. Lipopolysaccharide (LPS) detection by Toll-Like-Receptor 4 (TLR4) promotes DCs maturation by triggering a series of signaling cascades resulting in secretion of polarizing and inflammatory cytokines, upregulation of co-stimulatory molecules, as well as enhanced antigen processing and presentation (Mellman, 2013). All these functions are accompanied by major remodeling of membrane trafficking and actin organization to favor both antigen phagocytosis and DC migration to the lymph nodes (West et al., 2004, Bretou et al., 2017, Chabaud et al., 2015, Arguello et al., 2016). TLR4 is particularly expressed in the group 2 of conventional DCs (cDC2), characterized by the expression of surface CD11c, CD172a, as well as CD11b and endowed with a strong MHC II restricted antigen presentation capacity (Dalod et al., 2014).

One the major features of DC activation is a large increase in protein synthesis (Lelouard et al., 2007, Reverendo et al., 2019). This is required for the up-regulation of co-stimulatory molecules at the cell surface and acquire T cell immune-stimulatory function. Among the many pathways controlling mRNA translation, phosphorylation of eukaryotic initiation factor 2 (eIF2) is a central hub for regulating protein synthesis during stress. In homeostatic conditions, eIF2 mediates the assembly of the translation initiation complex and regulates start codon recognition in the mRNA by promoting the recruitment of other components necessary for active protein synthesis. During stress, phosphorylation of the alpha subunit of eIF2 (eIF2α) on serine 51 is mediated by a group of four eIF2α kinases (EIF2AK1-4), that specifically sense physiological imbalance (Claudio et al., 2013, Arguello et al., 2016). Phosphorylation of eIF2α converts eIF2 into a competitive inhibitor of the GDP-GTP guanidine exchange factor (GEF) eIF2B, impairing the GDP-GTP recycling required to form new translation initiation complexes and proceed with active mRNA translation (Yamasaki and Anderson, 2008). High levels of eIF2α phosphorylation impacts cells in two main ways: i) By reducing the rate of translation initiation and thus global protein synthesis levels; ii) By favoring the translation of the activating transcription factor 4 (ATF4) (Fusakio et al., 2016, Han et al., 2013), that in turn activates the transcription of many genes involved in the integrated stress response (ISR). The ISR aims at protecting cells from amino acid deprivation, oxidative, or mitochondrial stress and is incorporated as a branch of the ER unfolded protein response (UPR) upon PERK-activation. The ISR comprises a negative feedback loop that causes eIF2α dephosphorylation, through the induction of GADD34 (also known as PPP1R15a), a phosphatase 1 (PP1c) co-factor (Novoa et al., 2001, Harding et al., 2009). Consequently, dephosphorylation of p-eIF2α by GADD34/PP1c complexes, and associated protein synthesis restoration, signal ISR termination and a return to cellular homeostasis (Novoa et al., 2001). If stress persists, despite GADD34 activity, long term ATF4 expression promotes programmed cell death, through the induction of the pro-apoptotic transcription factor CHOP (Marciniak et al., 2004). Interestingly, ATF4 also regulates the expression of Rho GTPases and can control cell motility (Pasini et al., 2016), while globular actin has been shown to be part of the PP1c/GADD34 complex and provide additional targeting specificity for dephosphorylating p-eIF2α (Chambers et al., 2015, Chen et al., 2015).

The process of MAMPs sensing severely affects cellular homeostasis by boosting protein synthesis and results in a panoply of biochemical insults, that trigger various responses including autophagy and the ISR (Valecka et al., 2018, Arguello et al., 2016). the ISR can enter in a crosstalk with specialized microbe sensing pathways, which turns on or amplifies inflammatory cytokines production (Claudio et al., 2013, Reverendo et al., 2018). Toll-like receptor (TLR) activation in macrophages undergoing an ISR, suppress CHOP induction and protein synthesis inhibition, preventing apoptosis in activated cells (Woo et al., 2009). Moreover, ATF4 binds Interferon Regulatory factor-7 (IRF7) to block its capacity to induce transcription of type-I Interferon (IFN) induction (Liang et al., 2011). Several key innate immunity signaling cascades are believed to be dependent for their signalosome assembly on the chaperone HSPB8 and the kinase heme-regulated inhibitor (HRI/EIF2AK1) (Pierre, 2019). Microbe-activated HRI was shown to mediate phosphorylation of eIF2α and increase ATF4-dependent expression of HSPB8, thus promoting the signal transduction pathways leading to inflammatory cytokines transcription in macrophages (Abdel-Nour et al., 2019).

We show here that DCs obtained from spleen or derived from Fms-related tyrosine kinase 3 ligand (Flt3-L) treated-bone marrow cultures, display unusually high levels of phosphorylated eIF2α. Using Cre/*lox* recombination to generate mice specifically lacking GADD34 (PPP1R15a) or PERK (EIF2AK3) activity in DCs, we demonstrate that PERK-dependent eIF2α phosphorylation is acquired during DC development. PERK drives high eIF2α phosphorylation in steady state DCs with a relatively low impact on protein synthesis. We found that mRNA translation in DCs, differently to what has been shown during chronic ISR (Guan et al., 2017), is mediated despite high p-eIF2α levels by an eIF4F-dependent mechanism. We also found that primary DCs, conversely to macrophages, do not rely on eIF2α phosphorylation to activate cytokines transcription nor IL-1β secretion in response to MAMPs (Abdel-Nour et al., 2019),(Chiritoiu et al., 2019). Importantly, GADD34 antagonizes PERK activity to maintain functional protein synthesis levels in non-activated DCs. GADD34 upregulation in response to LPS, occurs in an ATF4-independent and TBK1-dependent manner, and contributes therefore directly to DC function by modulating the production of pro-inflammatory cytokines. Surprisingly, PERK activity impacts positively DC migration speed, correlating with the regulation of p-eIF2α levels by the synergistic action of GADD34 and actin cytoskeleton reorganization. Thus, DCs require PERK and GADD34 activity to coordinate protein synthesis, activation and migration capacity in response to MAMPs. These developmentally regulated features also endow DC with increase resistance to acute ER stress, preventing ATF4 induction in response to stressors like the bacterial subtilase cytotoxin (SubAB).

## Material and Methods

### Cell culture

Bone-marrow (BM) was collected from 6-9 weeks old female mice, and differentiated in DCs or macrophages during 7 days. The culture was kept at 37°C, with 5% CO_2_ in RPMI (GIBCO), 10% FCS (Sigma Aldrich), 100 units/ml penicillin, 100 units/ml streptomycin (GIBCO) and 50µM β-mercaptoethanol (VWR) supplemented with Flt3-L, produced using B16-Flt3-L hybridoma cells (Naik et al., 2005) for DC differentiation or M-CSF for macrophages, as described previously (Wang et al., 2013). For the migration assays, granulocyte-macrophage colony stimulating factor (GM-CSF) was used instead of Flt3-L and cells were cultured during 10-12 days with changes in the medium each 3 days. GM-CSF was obtained from transfected J558 cells (Pierre et al., 1997).To obtain splenocytes, spleens were collected, and injected with Liberase TL (Roche) and incubated 25 min at 37°C to disrupt the tissues. DC purification was performed using a CD11c+ positive selection kit (Milteny), according to manufacturer’s instructions and CD8α+ T cells isolation was performed with a Dynabeads™ untouched™ mouse CD8 T cells kit from Thermo Fischer Scientific. CD8α+ T where incubate overnight with anti-CD3 (10µg/ml) and anti-CD28 (5µg/ml) antibodies, to mimic activation by APCs. Mouse embryonic fibroblasts used in this work (MEFs), ATF4-/- and matched WT (129 SvEv) were a kind gift from Prof. David Ron (Cambridge Institute for Medical Research, UK). PERK KO-/- and matched WT were a kind gift from Prof. Douglas Cavener (Penn State University). MEFs were cultured in DMEM medium (GIBCO) with 5% FBS (Sigma) and 50 μM 2-mercaptoethanol. For the experimental assays, cells were plated from 16-24h before stimulation in 6 well plates, at 150 000 cells/ml in 2 ml of the same medium. After stimuli, cells were treated with trypsin-EDTA for 2 minutes at 37°C before washing in order to detach cells from the wells.

### Reagents

Lipopolysaccharide (Escherichia coli O55:B5), Cycloheximide, Puromycin, MRT67307 and Rocaglamide and Thapsigargin were purchased from Sigma Aldrich. Harringtonine is from ABCAM, Latrunculin A and Jasplakinolide are from Merck-Millipore. Low molecular weight Polyinosinic-polycytidylic acid (LMW poly(I:C)) was from InvivoGen. SAR1 was kindly provided by Sanofi (France) and Integrated Stress Response Inhibitor (ISRIB) was a gift from Carmela Sidrausky and Peter Walter, UCSF, San Francisco). Subtilase cytotoxin (Shiga toxigenic Escherichia coli strains) was purified from recombinant E. coli, as previously described (Paton et al., 2004). 4EGI-1 was purchased by Bertin bioreagent.

### Flow cytometry analysis

Cell suspensions were washed and incubated with a cocktail of coupled specific antibodies for cell surface markers in Flow activated cell sorting (FACS) buffer (PBS, 1% FCS and 2 mM EDTA) for 30 min at 4°C. For Flt3-l BMDCs the antibodies used were: CD11c (N418), SiglecH (551), CD86 (GL-1), F4/80 (BM8), CD64 (X54-5/7.1) from BioLegend; Sirpα (P84), CD24 (M1/69), MHCII (M5/114.15.2) from eBioscience™ CD11b (M1/70) from BDBioscience. For splenic cells, the antibodies used were NKp46 (29A1.4), CD4 (RM4-5), CD3 (145-2C11), CD11c (N418), CD19 (eBio1D3), CD8α (53-6.7) from eBioscience™, BST2 (927), Ly6G, F4/80, Ly6C from BioLegend; CD11b (M1/70), B220 (RA3-6B2) from BDBiosciences. These antibodies were used in combination with the LIVE/DEAD **®** Fixable Aqua Dead Cell Stain (ThermoFisher). For intracellular staining, cells were next fixed with BD phosflow fix buffer I (BD Biosciences) during 10 min at room temperature and washed with 10% Perm/wash Buffer I 1X (BD Biosciences). Permeabilized cells were blocked during 10’ with 10% Perm/wash buffer 1X, 10% FCS, before staining with primary antibodies. When the primary antibody was not coupled, cells were washed after and blocked during 10 min with Perm/wash buffer 1X, 10% FCS, 10% of serum from the species where the secondary antibody was produced. Then, the incubation with the secondary antibody was performed at 4°C during 30 min. p-eIF2α(S51) was purchased from ABCAM and p-eEF2(Thr56) from Cell signaling and Deoxyribonuclease (DNAse I) was purchased from Invitrogen. Data were acquired on an LSR-II/UV instrument using FACS Diva software. The acquired data were analyzed with FlowJo software (BD Biociences).

### Translation intensity and speed measurement

SUnSET technique to measure the intensity of protein synthesis was used as previously described (Schmidt et al., 2009). Puromycin was added in the culture medium at 12.5 μg/ml and the cells were incubated for 10 min at 37°C and 5% CO2 before harvesting. Cells were washed with PBS prior to cell lysis and immunoblotting with the anti-puromycin 12D10 antibody (Merck Millipore). For flow cytometry (flow) cells were processed, as described below for the intracellular staining, using the α-puromycin 12D10 antibody directly conjugated with Alexa 488 or A647 from Merck Millipore. The SUnRISE technique was performed as described (Arguello et al., 2018). Samples were treated with 2µg/ml of harringtonine at different time-points (90, 60, 45, 30, 15, 0 sec), and then treated with 12.5µg/ml of puromycin all at the same time. For the measurement of Cap-dependent translation cells were treated with 4EGI-I (100µM) or Rocaglamine (RocA-1) (100nM) for 0.5, 1, 2, or 4h. Cells were then incubated for 15 min at 37°C and stained with the 12D10 antibody (Merk-Millipore).

### Gene expression analysis

Total RNA was extracted from the DCs using the RNeasy mini kit (QIAGEN), including a DNA digestion step with RNAse-free DNAse (QIAGEN) and cDNA was synthesized using the Superscript II reverse transcriptase (Invitrogen). Quantitative PCR amplification was performed using SYBR Green PCR master mix (Takara) using 10ng of cDNA and 200nM of each specific primer on a 7500 Fast Real-PCR system (Applied Biosystems). cDNA concentration in each sample was normalized to GAPDH expression. The primers used for gene amplification were the following: GADD34 (S 5′-GACCCCTCCAACTCTCCTTC -3′, AS 5 ′-CTTCCTCAGCCTCAGCATTC -3′); IL-6 (S 5′-CATGTTCTCTGGGAAATCGTG -3′, AS 5 ′-TCCAGTTTGGTAGCATCCATC -3′); IFN-β (S 5′-CCCTATGGAGATGACGGAGA -3′, AS 5 ′-ACCCAGTGCTGGAGAAATTG -3′); IL-12 (S 5′-GGAATGTCTGCGTGCAAGCT-3′, AS 5 ′-ACATGCCCACTTGCTGCAT -3′); ATF4 (S 5′-AAGGAGGATGCCTTTTCCGGG -3′, AS 5 ′-ATTGGGTTCACTGTCTGAGGG -3′); CHOP (S 5′-CACTTCCGGAGAGACAGACAG -3′, AS 5 ′-ATGAAGGAGAAGGAGCAGGAG -3′); PERK (S 5′-CGGATTCATTGAAAGCACCT -3′, AS 5 ′-ACGCGATGGGAGTACAAAAC -3′); XBP1 (S 5’-CCGCAGCACTCAGACTATG -3’, AS 5’-GGGTCCAACTTGTCCAGAAT -3’); spliced XBP1 (S 5’-CTGAGTCCGCAGCAGGT -3’, AS 5’-AAACATGACAGGGTCCAACTT -3’); GAPDH (S 5′-TGGAGAAACCTGCCAAGTATG -3′, AS 5 ′-GTTGAAGTCGCAGGAGACAAC -3′); IL1-β (S 5’-TGATGTGCTGCTGCGAGAGATT -3’, AS 5’-TGCCATTTTGACAGTGA -3’); eIF2Bε (S 5’-GAGCCCTGGAGGAACACAGG -3’ AS 5’-CACCACGTTGTCCTCATGGC -3’); BIP S (5’ATTGGAGGTGGGCAAACCAA -3’ AS 5’-TCGCTGGGCATCATTGAAGT -3’).

### Gene set enrichment analysis

The gene set enrichment analysis has been performed using published murine microarray datasets accessible through the Gene Expression Omnibus (GEO) repository under the references GSE9810 (Robbins et al., 2008) and GSE2389 (Fontenot et al., 2005). Raw microarray data describing CD8∝ + cDCs (DC1), CD11b+ cDC (DC2), pDC (plasmacytoid dendritic cells), B cells, NK cells, CD8+ T cells (Robbins et al., 2008) and CD4+ T cells (Fontenot et al., 2005) in mice spleen have been download For each of these cell types, the hybridation has been performed using Affymetrix mouse 4302.0 gene chips. Microarray data have been normalized by Robust Multi-array Average (RMA) algorithm (Irizarry et al., 2003) using the oligo BioconductorR package (Carvalho and Irizarry, 2010). Normalization consists of a background correction of raw intensities, a log2 transformation followed by quantile normalization in order to allow the comparison of each probe for each array. Prior data usage, the absence of batch effect was assessed by Principal Component Analysis (PCA) using ade4R package (Bougeard and Dray, 2018).

Gene set enrichment analysis (GSEA) was performed using publicly available gene signatures reflecting an ISR state (Supplementary Table 1, 2 and 3). Lists of ATF4 and CHOP target genes identified by ChIP-seq experiments (Han et al., 2013) have been used to search for ATF4-dependent and CHOP-dependent signatures in DCs (Supplementary table 2 and 3). Lists of genes for which a translational upregulation after 1h of thapsigargin (Tg) treatment and congruent (both transcriptional and translational) upregulations after 16h of Tg treatment (Guan et al., 2017) were used to search gene expression signatures respectively of acute and/or chronic ISRs in DCs. GSEA was generated using BubbleGUM (Spinelli et al., 2015). Briefly, GSEA pairwise comparisons are performed for each probe and the multiple testing effects are corrected using a Benjamini-Yekutieli procedure. The corrected p-values are hence calculated based on a null hypothesis distribution built from the permutations of the gene sets across all the pairwise comparisons. In our analyses,10,000 permutations of the gene sets have been performed in order to compute the p-values. All results with a false discovery rate (FDR) below the threshold of 0.25 have been considered as significant.

### Immunoblotting

Cells were lysed in RIPA buffer (25mM Tris-HCl pH 7.6, 150mM NaCl, 1% NP-40; 1% sodium deoxycholate, 0.1% SDS) supplemented with Complete Mini Protease Inhibitor Mixture Tablets (Roche), NaF (Ser/Thr and acidic phosphatase inhibitor), NaVO4 (Tyr and alkaline phosphatase inhibitor) and MG132 (proteasome inhibitor). The nuclear extraction was performed using the Nuclear Extract kit (Active Motif) according with manufacturer’s instructions. Protein quantification was performed using the BCA Protein Assay (Pierce). Around 20 µg of soluble proteins were run in 4-20% acrylamide gradient gels and for the immunoblot the concentration and time of incubation had to be optimized for each individual antibody. Rabbit antibodies against eIF2α, p-eEF2(Thr56), eEF2, eIF2B, p-IRF3 (ser396), IRF3, p-S6, PERK, were purchased from Cell Signaling (ref 5324, 2331, 2332, 3592, 4947, 4302, 2211, 3192 respectively). Rabbit antibody against p-eIF2α(S51) was purchased from ABCAM (Ref 32157). Rabbit antibody against ATF4 was purchased from Santa Cruz Biotechnology (sc-200). Mouse antibody against β-actin was purchased from Sigma (A2228). Mouse antibodies against HDAC1 and S6 were purchased from Cell Signaling (ref 5356, 2317 respectively). Mouse antibody against puromycin was purchased from Merck Millipore (MABE343). Mouse antibody against p-eIF2β was a kind gift from David Litchfield (University of Western Ontario, Ca). Mouse antibody against eIF2β was purchased from Santa Cruz Biotechnology (sc-9978). HRP Secondary antibodies were from Jackson ImmunoResearch Laboratories.

### Cytokines measurement

The IL-6, the IFN-β, and IL1β quantifications from the cell culture supernatant were performed using the mouse Interleukin-6 ELISA kit (eBioscience), the mouse IFN-β ELISA kits (PBL-Interferon Source or Thermos Fisher) according to manufacturer’s instructions.

### Immunohistochemistry

Spleens were snap frozen in Tissue Tek (Sakura Finetek). Frozen sections (8 μm) were fixed with acetone permeabilized with 0,05% saponin. The following antibodies were used for the staining: CD11c (N418) from Biolegend (Ref 117301), p-eIF2α (Ser 52) from Invitrogen (Ref 44-728G), CD11b (M1/70) from BD Biosciences, CD8α-biotin (53-6.7) Biolegend (Ref 100703) and B220 (RA3-6B2) from Invitrogen (Ref 14-0452-81). Images were collected using a Zeiss LSM 510 confocal microscope. Image processing was performed with Zeiss LSM software.

### Mice

Wild-type (WT) female C57BL/6 mice were purchased from Janvier, France. PKR-/- C57BL/6 were a kind gift from Dr. Bryan Williams (Hudson Institute of Medical Research, Australia) (Kumar et al., 1997). PERK^loxp/loxp^ mice were the kind gift of Dr Doug Cavener (Zhang et al., 2006) and purchased from Jackson Laboratories. GADD34ΔC^loxp/loxp^ mice were developed at the Centre d’Immunophénomique (CIPHE, Marseille, France). PERK^loxp/loxp^ and GADD34^loxp/loxp^ were crossed with Itgax-Cre+ mice (Caton et al., 2007) and backcrossed, to obtain stable homozygotic lines for the loxp sites expressing Cre. For all studies, age-matched WT and transgenic 6 to 9 weeks females were used. All animals were maintained in the animal facility of CIML or CIPHE under specific pathogen–free conditions accredited by the French Ministry of Agriculture to perform experiments on live mice. These studies were carried out in strict accordance with Guide for the Care and Use of Laboratory Animals of the European Union.

All experiments were approved by the Comité d’Ethique PACA and MESRI (approval number APAFIS#10010-201902071610358). All efforts were made to minimize animal suffering.

### Preparation of microchannels and speed of migration measurement

Microchannels were prepared as previously described (Vargas et al., 2016). For velocity measurements (carried out in 4µm-by-5µm microchannels), phase contrast images of migrating cells were acquired during 16 hours (frame rate of 2 min) on an epifluorescence Nikon Ti-E video microscope equipped with a cooled charge-coupled device (CCD) camera (HQ2, Photometrics) and a 10× objective. Kymographs of migrating cells were generated and analyzed using a custom program.

### Statistical analysis

Statistical analysis was performed using GraphPad Prism Software. The most appropriate statistical teste was chosen according to the set of data. Mainly, we used Student’s t test and Kruskal-Wallis test were used. * *p* < 0.5, ** *p* <0.01, *** *p* < 0.001, **** *p* < 0.0001.

## Results

### Steady-state dendritic cells display high levels of eIF2α phosphorylation

Given the importance of eIF2α in cell homeostasis, we monitored the physiological levels of phosphorylated eIF2α (p-eIF2α) in mouse spleen sections by immunohistochemistry. All DC subsets, identified by virtue of CD11c expression together with CD8α (cDC1), CD11b (cDC2) or B220 (pDC), displayed high levels of eIF2α phosphorylation (Fig. 1A and 1B), strongly contrasting with other splenocytes, like B cells (Fig. 1B lower panel). Splenocytes isolation and immunodetection based-quantification of eIF2α phosphorylation confirmed the high levels of p-eIF2α observed in different DC subsets compared to other cells including macrophages (F4/80+), T cells (CD3+/CD4+ or /CD8+) and B cells (Fig. 1C). We next evaluated the levels of p-eIF2α in BM-derived DCs differentiated in presence of Flt3-Ligand (Flt3-L BMDC), encompassing the major cDC1, cDC2 and pDC subsets in different proportions (circa 30%, 60% and 10% respectively of the CD11c+ population) and with phenotypes equivalent to those of spleen DC subsets (Brasel et al., 2000). This was further confirmed upon sorting and analysis of the different populations, which were submitted to p-eIF2α immunoblot in comparison to isolated primary CD8+ T cells or mouse embryonic fibroblasts (MEFs) stimulated or not with the ER-stress inducing drug thapsigargin for 2h (Fig. 1D). eIF2α phosphorylation was evident in all DC subsets, even when compared to thapsigargin-treated control T cells and MEFs. Quantification of p-eIF2α/eIF2α ratios indicated that steady state DCs display 2 to 4 times more p-eIF2α, than stressed MEFs, with the cDC1 population displaying the highest ratio of phosphorylation (Fig. 1D). We next evaluated when eIF2α phosphorylation was acquired during DC differentiation. The daily immunoblot analysis of differentiating Flt3-L BMDCs, allowed us to establish that high p-eIF2α levels were displayed by DCs from 4 days of culture (Fig. 1E). This suggests that this event is an integral part of the DC development program induced by Flt3-L (Fig. 1E). Given the described dominant negative effect of p-eIF2α on translation initiation, we next sought to monitor the levels of protein synthesis in the different DC subsets. CD8+ T cells were used as a reference, since in these cells, p-eIF2α is barely detectable. We used puromycilation and detection by flow cytometry (flow) (Schmidt et al., 2009, Arguello et al., 2018) to establish that, despite high eIF2α phosphorylation levels, all resting DC subsets displayed higher level of translation compared with resting and CD3/CD28 stimulated CD8+ T cells (Fig. 1F). We next monitored protein synthesis every two days of culture to establish precisely the influence of eIF2α phosphorylation over protein synthesis activity during DC differentiation *in vitro*. After puromycilation and flow analysis using various DC differentiation markers, we applied dimensionality reduction using t-distributed stochastic neighbor embedding (tSNE) to visualize DC differentiation and the evolution of protein synthesis activity within the different subpopulations over time (Fig. 2A). We confirmed that protein synthesis levels steadily increased with differentiation in all three DC subsets, this despite high eIF2α phosphorylation. These observations suggest that steady state DCs have adapted their translation machinery to overcome the dominant negative effect on translation initiation of eIF2α phosphorylation on Ser51, which is clearly associated with their differentiation from bone marrow progenitors.

**Figure 1.**
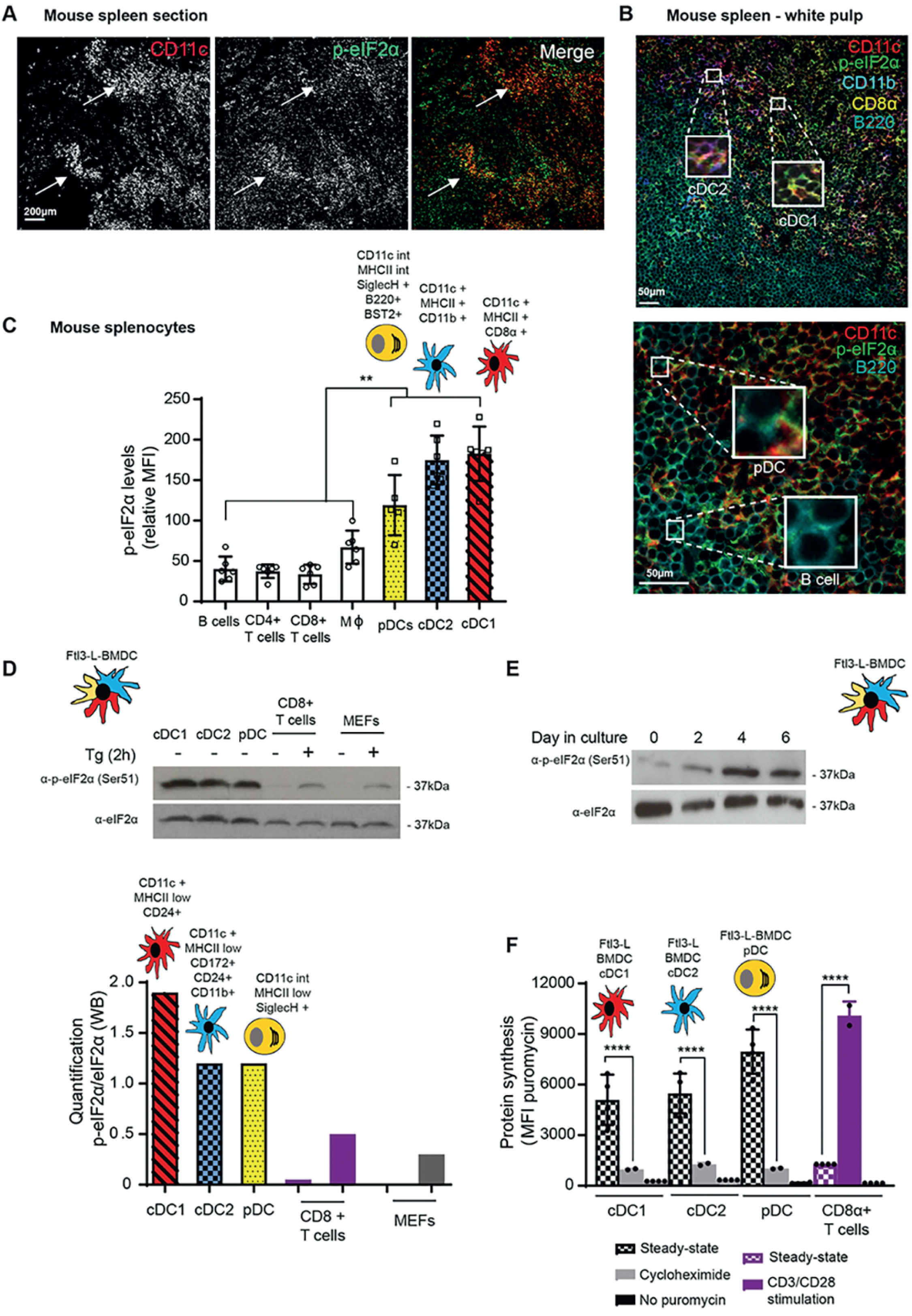
Steady-state Flt3-L BMDCs and splenic DCs display remarkably high levels of eIF2α without inhibition of translation. (A) Immunohistochemistry of mouse spleen with staining for CD11c (red) and p-eIF2α (green). scale bar 200µm, magnification 10X. Single color images are shown and merged picture (right row) indicates that high level of p-eIF2α staining is mostly found co-localizing in cells positive for CD11c+ (DCs, white arrowheads). (B) Immunohistochemistry of mouse spleen in the white pulp for CD11c (red), p-eIF2α (green), CD11b (blue), CD8α (yellow) and B220 (turquoise), scale bars 50µm, magnification 40x. In the upper panel, Magnified areas shows p-eIF2 detection in cDC2 (CD11c+/CD11b+, purple) and cDC1 (CD11c+/CD8α+, orange). In the lower panel, magnified areas show p-eIF2 detection in pDCs (B220+/CD11c+) and in B cells (B220+ and CD11c-). (C) Relative p-eIF2α levels measured by flow in different mouse spleen populations. DC subsets display higher levels of p-eIF2α in comparison to other splenocytes populations. Statistic analysis was performed by Kruskal-Wallis test, followed by paired pos-hoc test corrected with Benjamin Holscher procedure. ** *p* < 0.01 (D) Levels of p-eIF2α and total eIF2α were measured in DC populations by immunoblot. Sorted steady-state Flt3-L BMDCs were compared with MEFs and freshly isolated CD8α+ T cells stimulated or not with thapsigargin (Tg) for 2h (200nM). Ratio of p-eIF2α/eIF2α is quantified in the graph of the lower pannel. (E) Levels of p-eIF2α and total eIF2α were measured in bulk Flt3-L BMDCs during different days of bone marrow differentiation *in vitro*. (F) levels of protein synthesis were measured by puromycilation and intracellular flow cytometry detection in different subsets of Flt3-L BMDCs and in CD8+ splenic T cells. Cells were incubated with puromycin 10 min before harvesting and when indicated, cycloheximide (CHX, 10µM) was added 5 minutes before puromycin. Steady-state Flt3-L BMDCs were directly compared with CD8+ splenic T cells either steady-state or stimulated overnight with anti-CD3 (10ug/ml) and anti CD28 (5ug/ml). Samples without previous incorporation of puromycin were used as control. All data are representative of n=3 independent experiments. Data in (F) represent MFI +/- SD of 3 independent experiments. Unpaired Student’s t-test (**** *p* < 0.0001).

**Figure 2.**
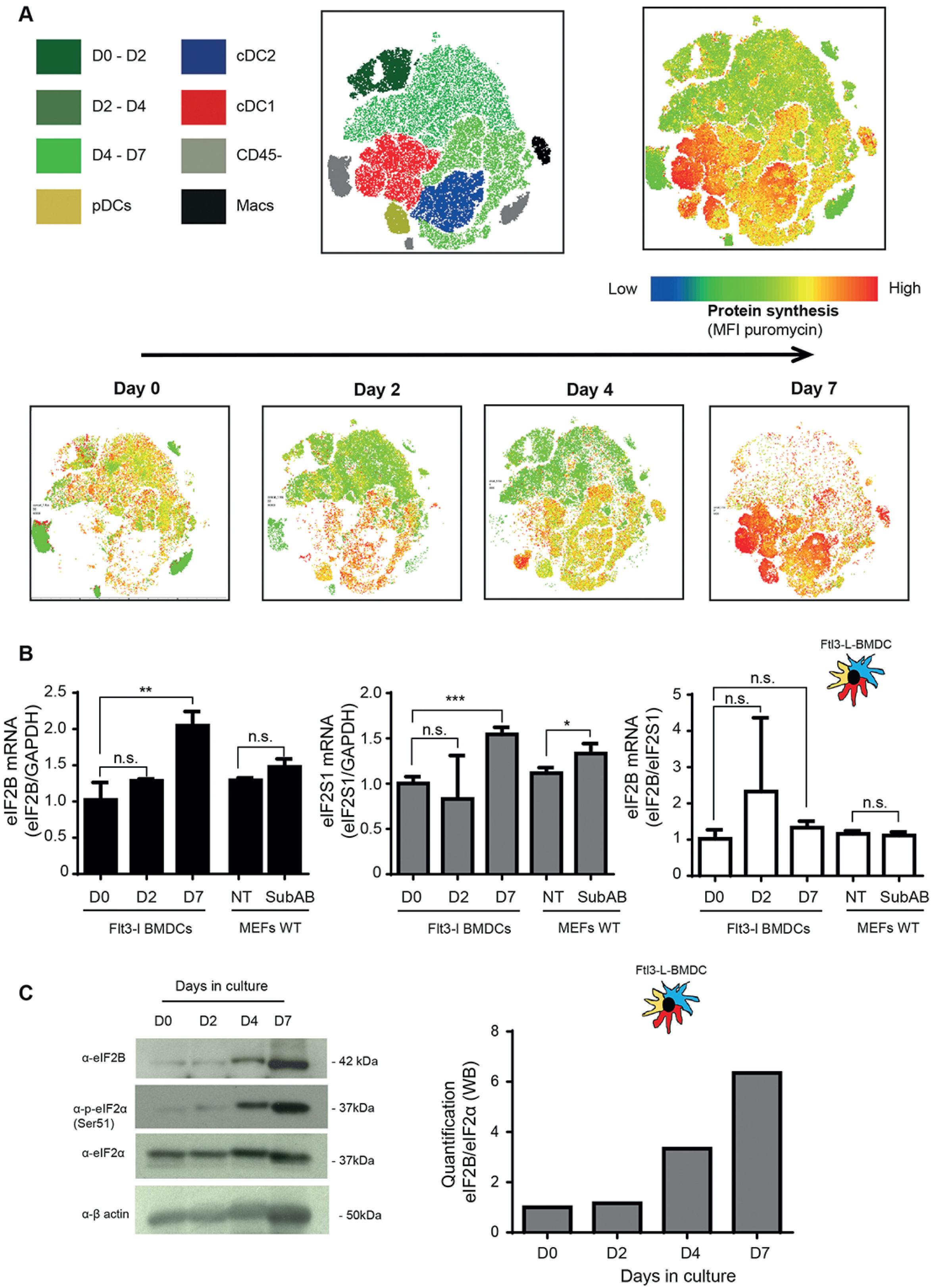
Protein synthesis is increased during in BMDCs differentiation. A) Levels of protein synthesis were measured every two days by flow cytometry during Flt3-L BMDCs differentiation *in vitro* (0, 2, 4 and 7d). Cells were incubated during 10 min with puromycin, intracellularly stained with an α-puromycin antibody prior analysis. The same dimensionality reduction using t-distributed stochastic neighbor embeding were applied to all samples. Macrophages (Macs) in black are gatted as CD45+, CD11c+, CD11b+, F4/80+, CD64+ cells; cDC1 in red express CD45+, CD11c+, MCHII+, CD24+; cDC2 in red express CD45+, CD11c+, MHCII+, CD11b+, Sirpα+, pDC in yellow express CD11c^int^ and Siglec H+, cells negative for CD45 in grey are considered as non immune, and progenitors are stain in green. B) mRNA levels of eIF2B*ε* and eIF2α (eIF2S1) measured by RT-qPCR in bulk Flt3-L BMDCs at indicated days of differentiation and compared with control MEFs (WT) at steady state or treated during 1h with SubAB (250ng/ml). Quantification of the ratio eIF2B/eIF2S1 in shown on the right panel. C) Levels of eIF2B*ε*, p-eIF2α total eIF2α and β-actin were measured in bulk Flt3-L BMDCs at indicated days of differentiation *in vitro*. In parallel to eIF2 α phosphorylation, eIF2B levels are strongly up-regulated during DC differentiation. Quantification of the ratio eIF2B/eIF2α is shown on the right. All data are representative of n=3 independent experiments. Data in (B) represent Mean +/- SD of 3 independent experiments. Unpaired Student’s t-test (* *p* < 0.05, ** *p* < 0.01 and *** *p* < 0.001).

### eIF2B expression is strongly up-regulated upon DC differentiation

P-eIF2α inhibits translation initiation by forming a stable inhibitory complex with eIF2B, that reduces the GEF activity of eIF2B. eIF2B levels are generally lower than those of eIF2 and partial eIF2α phosphorylation is sufficient to attenuate protein synthesis initiation in most cells (Adomavicius et al., 2019). We therefore monitored eIF2B*ε* expression levels during DC differentiation and observed that eIF2B*ε* levels were strongly increased both transcriptionally and translationally in differentiated cells (Fig. 2B & 2C). The progression of the ratio eIF2B*ε* over eIF2α proteins during differentiation (Fig. 2C), suggests that the quantity of p-eIF2α required to inhibit translation initiation in DCs is superior to what is needed in other cells like MEFs. This developmentally regulated expression of eIF2B and eIF2S1, could therefore explain the adaptation of DCs to relatively high p-eIF2α levels and the progressive acquisition of high translation levels upon differentiation.

### Flt3-L BMDCs activation by LPS promotes eIF2α-dephosphorylation

Phosphorylation of eIF2α and of elongation factor 2 (eEF2) were gradually lost during *E. coli* lipopolysaccharide (LPS)-stimulation of Flt3-L BMDCs, indicating a tight regulation of this pathway during DC activation (Fig. 3A). Complementary to p-eIF2α, phosphorylated eEF2 is a major repressor of translation in adverse growth conditions, such as starvation, or accumulation of misfolded proteins in the endoplasmic reticulum (ER) (Ryazanov, 2002), (Lazarus et al., 2017). Stress activates elongation factor 2 kinase (eEF2K), which phosphorylates eEF2 and inhibits translation elongation through an arrest of codon translocation from ribosomal A to P sites (De Gassart and Martinon, 2017, Richter and Coller, 2015). Similarly to eIF2α, eEF2 phosphorylation levels were found high in steady-state Flt3-L BMDCs and down-regulated after LPS stimulation (Fig. 3B). Given their sensitivity to LPS-activation, we monitored more specifically TLR4 expressing DC2 by flow, and observed a profound loss of p-eEF2 after 30 min of stimulation, thus demonstrating the strong impact of LPS signaling on this major translation factor and on protein synthesis intensity, which was doubled over 6h of stimulation (Fig. 3C).

**Figure 3.**
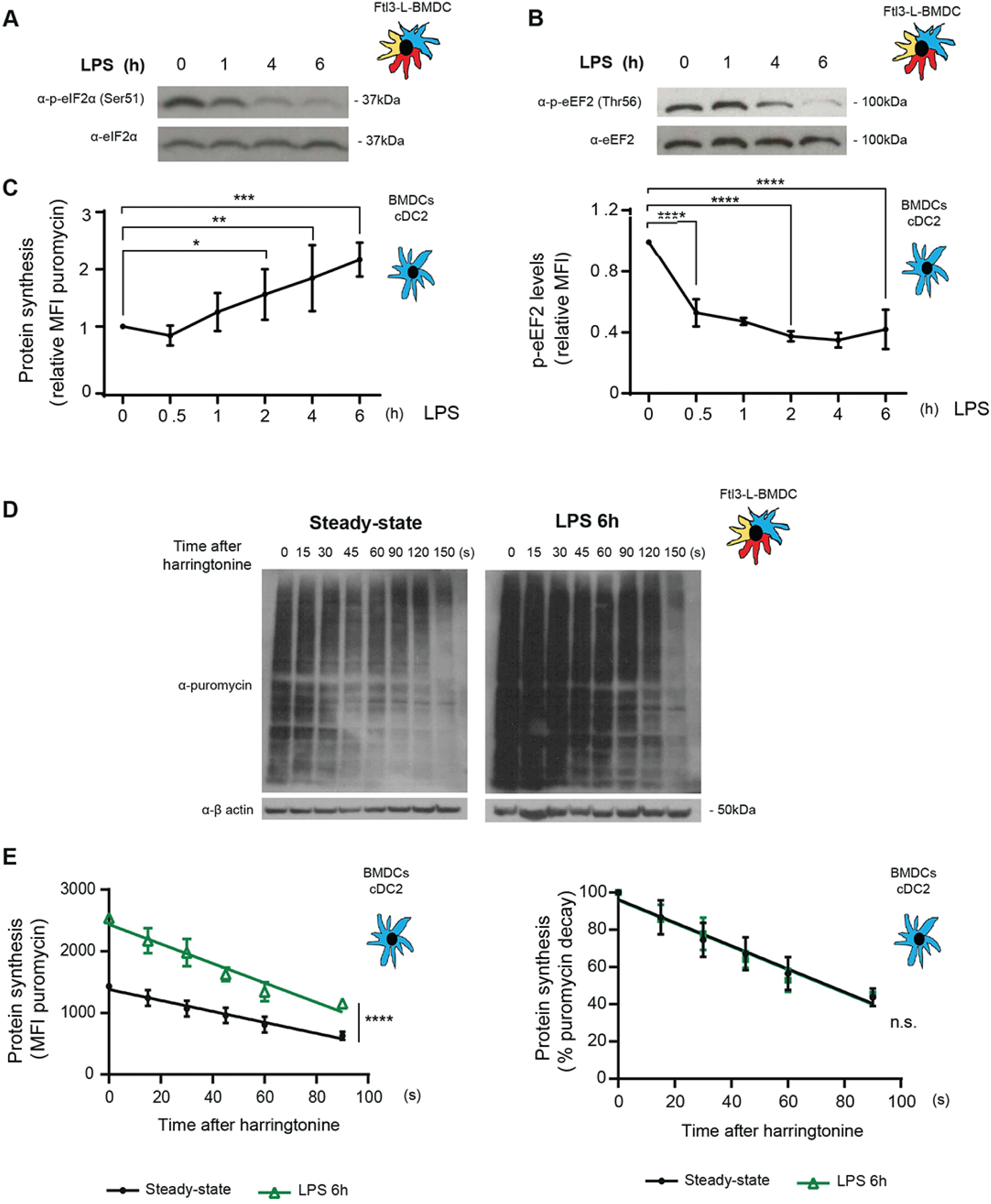
P-eIF2α and p-eEF2 levels are down-regulated upon LPS stimulation. Flt3-L BMDCs were stimulated with LPS (100ng/ml) for indicated hours. (A) Immunoblot for p-eIF2α and total eIF2α in bulk Flt3-L BMDCs. (B) Immunoblot for p-eEF2 (B) and total eEF2 in bulk Flt3-L BMDCs. Phosphorylation of both translation factors is reduced upon LPS activation. C) Monitoring of protein synthesis (puromycin) and p-eEF2 by intracellular flow in cDC2 (CD11c+, Siglec H-, CD172α+, CD24+). (D) and (E) Translation initiation and speed of elongation was measured using SunRISE after 6h of incubation with LPS. Flt3-L BMDCs were treated with harringtonine (2μg/ml) for different times up to 150 seconds prior incubation with puromycin during 10 min. (D) DC samples were lysed and subjected to immunoblot using an α-puromycin antibody and β-actin as loading control. (E) Flow cytometry detection of puromycin (SunRISE) gated on the cDC2 population. Left, total puromycin mean fluorescence intensity (MFI) decay between steady-state and LPS activated cells. Right, relative MFI decay, showing that the speed of elongation does change upon activation. All data are representative of n=3 independent experiments. Data in Figures (C) and (D) represent Mean +/- SD of 3 independent experiments. Unpaired Student’s t-test (* *p* < 0.05, ** *p* < 0.01, *** *p* < 0.001 and **** *p* < 0.0001).

Given the rapidity and intensity of eIF2α and eEF2 dephosphorylation upon activation, we applied to cDC2, the SunRISE technique, a method for monitoring the translation level and elongation rate of multiple cell types in parallel (Arguello et al., 2018). SunRiSE was applied using detection by immunoblot (Fig. 3D) and by flow (Fig. 3E), confirming a striking augmentation of translation intensity in LPS-activated DCs compared to the steady-state situation (T=0s). Polysomes elongation speed, indicated by the rate of puromycin staining decay after harringtonine treatment, remained however identical among steady-state and LPS-activated cDC2 (Fig. 3E). eIF2α and eEF2 dephosphorylation in activated DCs is therefore correlated with a strong increase in translation initiation, rather than elongation, to allow mRNA translation to reach its maximum level concomitantly with the acquisition of DC immune-stimulatory capacities (Lelouard et al., 2007).

### PPP1R15a (GADD34) controls eIF2α dephosphorylation in activated DCs

The inducible PP1c co-factor PPP1R15a, known as GADD34, is key in mediating p-eIF2α dephosphorylation in the resolution phase of the ISR (Novoa et al., 2001, Novoa et al., 2003). GADD34 is also absolutely necessary to re-establish protein synthesis in MEFs undergoing PKR activation upon dsRNA exposure or viral infection (Clavarino et al., 2012a). We have also shown that while induction of GADD34 transcription during ER stress is ATF4-dependent (Walter and Ron, 2011), expression of GADD34 upon viral sensors activation is Interferon Regulatory Factor 3 (IRF3 dependent (Dalet et al., 2017). We therefore investigated if the p-eIF2α dephosphorylation observed in Flt3-L BMDCs was GADD34 dependent, and whether TANK-binding kinase 1 (TBK1)/IRF3 signaling cascade, which is triggered upon LPS stimulation, was responsible for its transcriptional induction during DC activation. The IKKϵ/TBK1 inhibitor (MRT67307, TBKin) (Clark et al., 2011) was used to follow the impact of IKKϵ/TBK1/IRF3 activation on LPS-induced p-eIF2α dephosphorylation (Fig. 4A). Upon efficient treatment with TBKin, indicated by the loss of p-IRF3 detection, p-eIF2α levels were clearly augmented even in absence of LPS stimulation, suggesting that baseline IKKϵ/TBK1/IRF3 signaling decreases p-eIF2α levels by driving steady state GADD34 expression. LPS-dependent induction of GADD34 mRNA was also severely decreased upon IKKϵ/TBK1 inhibition (Fig. 4B), suggesting that in DCs, *Ppp1r15a/GADD34* transcription is partially dependent on IRF3 activation and not only on the ATF4-dependent transcriptional axis. This hypothesis was reinforced by the observation that ATF4 and CHOP expression, the two main transcription factors controlling *Ppp1r15a/GADD34* transcription during ER-stress, remained stable after 6h of LPS stimulation, and in the case of CHOP even decreased in the first hours of activation (Fig. 4C). Surprisingly, IKKϵ/TBK1 inhibition also augmented considerably MHC II surface expression (Fig. 4D), presumably by changing the levels of secreted cytokines influencing DC maturation in the culture. This was further exemplified by the absence of IRF3-dependent IFN-β expression in TBKin-treated cells (Fig. 4E), while NF-κB-dependent transcription of IL-6 remained unaffected (Fig. 4E), as well as CD86 surface appearance (Fig. 4D). Importantly, despite augmenting p-eIF2α levels, IKKϵ/TBK1 inhibition also strongly boosted protein synthesis suggesting that DCs conserve the capacity to augment their protein synthesis irrespective of TBK activation and despite a low rate eIF2α dephosphorylation (Fig. 4F). Artificial inhibition of IKKϵ/TBK1, prevents therefore GADD34 and IFN-β expression, while augmenting eIF2α phosphorylation. However it reveals also the existence of a signaling pathway downstream of TLR4, capable of overcoming augmented levels of p-eIF2α and of increasing protein synthesis, independently of GADD34 expression. Given the amount of eIF2B expressed by differentiated DCs (Fig. 2B and 2C), this pathway could be equivalent to the TLR-dependent activation of the eIF2B GEF through PP2A-mediated serine dephosphorylation of the eIF2Bϵ-subunit, that was observed by Woo and colleagues in macrophages (Woo et al., 2012).

**Figure 4.**
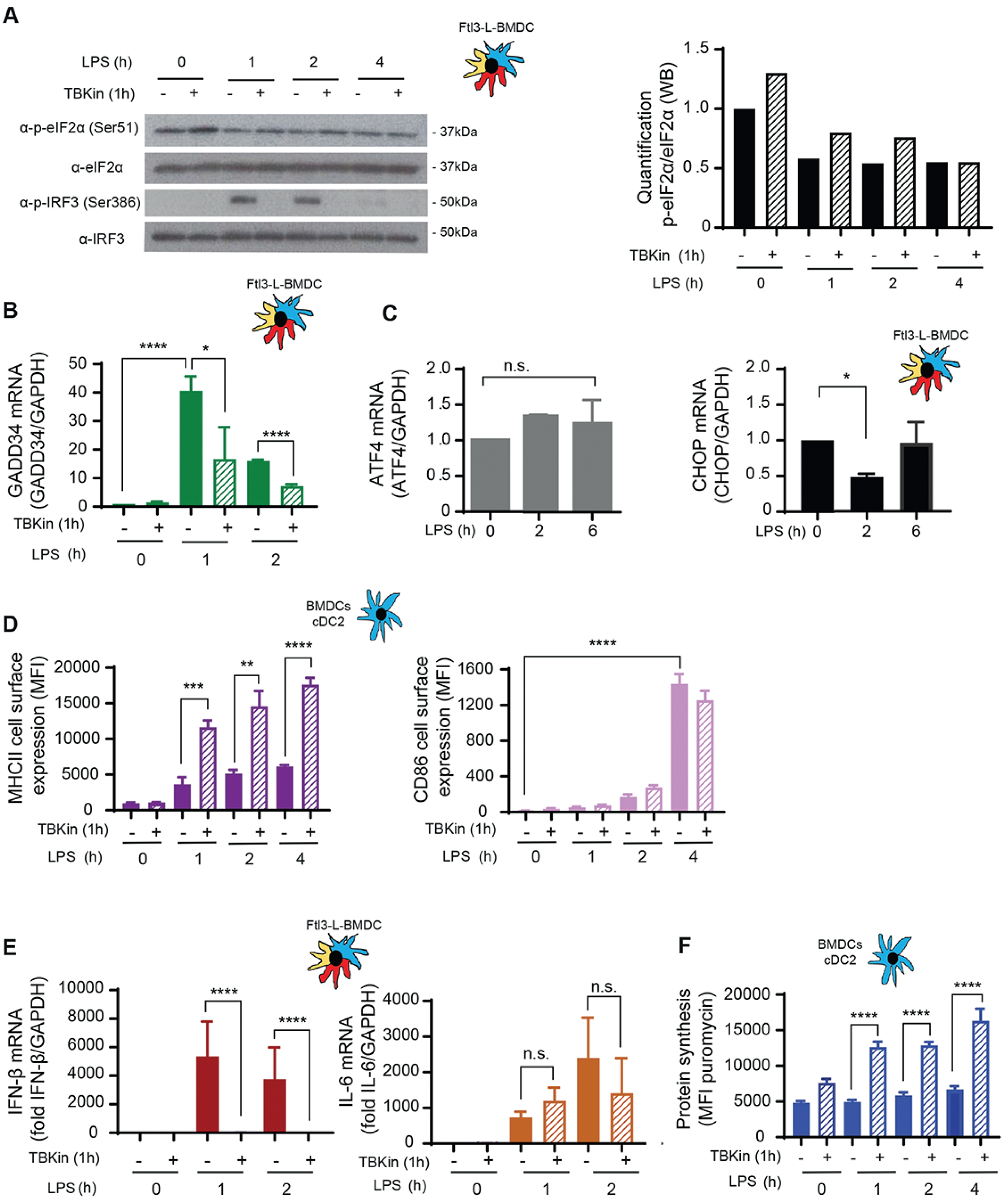
GADD34 expression is regulated by TBK1 signaling. Flt3-L BMDCs were pre-treated with 2µM of the TBK1 inhibitor (MRT67307) for 1h, before stimulation with LPS (100ng/ml) for indicated times. (A) Immunoblot detection of p-eIF2α, total eIF2α, P-IRF3 and total IRF3. Quantification is represented on the right. (B-E) mRNA levels of GADD34 (B), ATF4 (C), CHOP (C), IFN-β (E) and IL-6 (E) were measured by RT-qPCR in Flt3-L BMDCs. Data were normalized to the housekeeping gene (GAPDH) level. (D) Upregulation of co-stimulatory molecules CD86 and of MHC II was measured by flow cytometry in the cDC2 population (MFI). (F) Levels of protein synthesis were measured by puromycilation and flow in the cDC2 population from Flt3-L BMDCs. Cells were incubated with puromycin 10 min before harvesting. All data are representative of n=3 independent experiments. Data in Figures (B-F) represent Mean +/- SD of 3 independent experiments. Unpaired Student’s t-test (* *p* < 0.05, ** *p* < 0.01, *** *p* < 0.001 and **** *p* < 0.0001).

We further investigated the impact of GADD34 on the eIF2α pathway in DCs by generating, a novel transgenic mouse model with floxed alleles for *PPP1R15a*, allowing the deletion of the third exon coding for the C-terminal PP1 interacting domain, which upon Cre recombinase expression create a null phenotype for GADD34-dependent eIF2α dephosphorylation (Harding et al., 2009) (Sup Fig. S1A). *Ppp1r15a*^loxp/loxp^ C57/BL6 mouse were crossed with a Itgax-cre deleter strain (Caton et al., 2007) to specifically inactivate *ppp1r15a* activity in CD11c expressing cells, including all DC subsets. Despite inducing a light splenomegaly, *ppp1r15a* inactivation had no obvious consequences for splenocytes development *in vitro and in vivo* (Sup. Fig. S2). Flt3-L BMDCs derived from WT and Itgax-cre/*Ppp1r15a*^loxp/loxp^ (GADD34ΔC) mice were LPS activated prior detection of translation factors by immunoblot (Fig. 5A). GADD34 inactivation prevented LPS-dependent eIF2α dephosphorylation, however no impact was observed on the phosphorylation levels of the activator beta subunit of eIF2 (eIF2β), eEF2, nor ribosomal S6 protein, demonstrating that among these translation factors subunits, GADD34 only targets eIF2α with high specificity (Fig. 5A). Consequently, p-eIF2α levels do not influence directly the fate of the eIF2β subunit and eEF2 in DC, nor their respective impact on translation. eIF2β phosphorylation has been previously shown to counteract p-eIF2α dominant negative effect and promote mRNA translation (Gandin et al., 2016). However, it was neither impacted by LPS activation nor GADD34 activity in DCs, and thus unlikely to interfere with eIF2α regulation. Functional deletion of GADD34 inhibited translation initiation in both steady-state and LPS-activated cDC2 (Fig. 5B), and also reduced moderately translating polysomes speed in non-stimulated cells. GADD34 expression seems therefore to prevent exaggerated protein synthesis inhibition linked to abundant eIF2α phosphorylation in steady-state DCs. The amount of eIF2B present in the DC seems, however, sufficient to maintain a lower but still overall active protein synthesis despite GADD34 inactivation and increased P-eIF2α, as already suggested by our results obtained with TBKin treatment (Fig. 4F). These results confirm that GADD34 is produced and active in steady-state DC, suggesting the existence of a basal IRF3-dependent *Ppp1r15a* mRNA transcription in DCs. Interestingly these results also imply that GADD34 mRNA translation occurs independently of any acute stress in DCs, despite its upstream uORF-dependent translational regulation, as phenomenon recently observed in MEFs (Reid et al., 2016).

**Figure 5.**
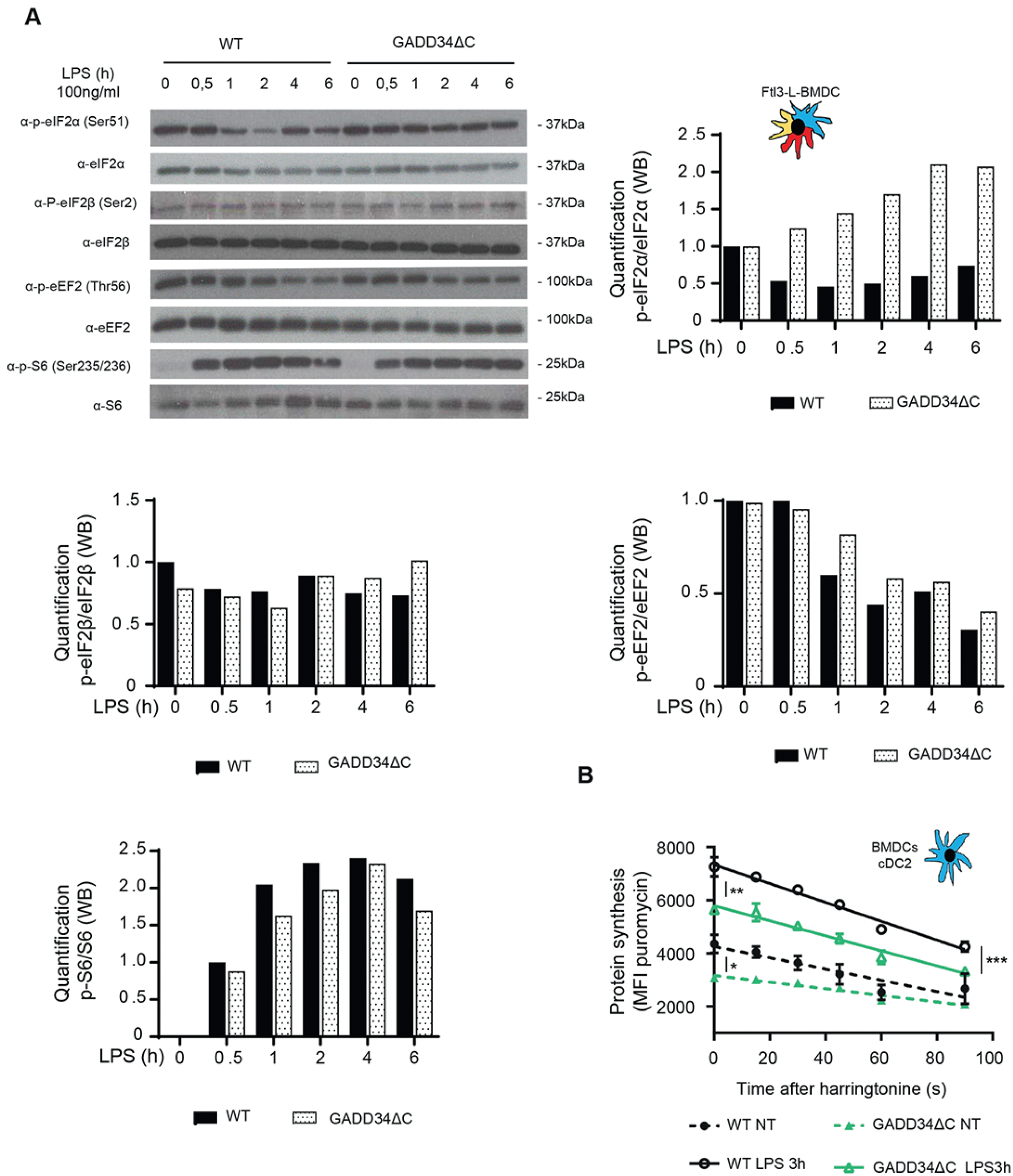
GADD34 mediates eIF2α dephosphorylation and controls translation in DCs. WT and GADD34ΔC BMDCs were stimulated with LPS (100ng/ml) for the indicated times. (A) Levels of p-eIF2α, total eIF2α, p-eIF2β, total eIF2β, p-eEF2, total eEF2, P-S6 and total S6 were detected by immunoblot (top left) and quantification is shown in the different panels. (B) The speed of translation elongation was measured by SunRISE after 3h of incubation with LPS. Harringtonine (2μg/ml) was added at different times up to 90 seconds prior incubation with puromycin during 10 min. Flow intracellular staining was performed in cDC2 using an α-puromycin antibody. The total decay of puromycin MFI between WT and GADD34ΔC in steady-state and upon activation indicates that translation initiation and elongation speed is decreased in GADD34-deficient DCs. All data are representative of n=3 independent experiments. Data in (B) represents MFI +/- SD of 3 independent experiments. Unpaired Student’s t-test (** *p* < 0.01).

### PERK mediates eIF2α phosphorylation in steady state DCs

We next investigated the consequences of inactivating known eIF2A kinases on the p-eIF2α levels observed in steady-state Flt3-L BMDCs (Krishna and Kumar, 2018). We tested pharmacological and genetic inactivation of PKR (EIF2AK2) and GCN2 (EIF2AK4) (Sup Fig. S3), without observing any major disturbances in eIF2α phosphorylation levels. We next turned towards the ER-stress kinase PERK (EIF2AK3) by crossing PERK^loxp/loxp^ mice with the Itgax-cre strain (Caton et al., 2007) allowing for the deletion of the exons 7-9, coding for the kinase domain (PERKΔK), and consequently PERK inactivation in CD11c-expressing cells (Sup Fig. S1B). PERK protein levels were enriched in WT CD11c+ splenic DC compared to other splenocytes (Fig. 6A), further suggesting the importance of this molecule for DC function. PERK expression was efficiently abrogated in CD11c+ splenocytes and Flt3-L BMDCs derived from animals bearing the floxed-PERK alleles (Fig. 6A), with no obvious consequences for splenocytes development *in vitro and in vivo* (Sup. Fig. S4). PERK inactivation decreased p-eIF2α levels by 60% in steady-state DCs (Fig. 6B), while p-eEF2 levels remained unchanged (Fig. 6C). Interestingly, LPS stimulation increased PERK levels in WT Flt3-L BMDCs, suggesting that this kinase could be further activated during DC maturation. However, despite increased PERK levels, p-eIF2α levels were nevertheless decreased with activation, reaching a minimum at 4h of LPS stimulation (Fig. 6B). This low phosphorylation intensity of eIF2α was equivalent to the basal levels observed in PERK-deficient DCs, and likely resulting from other eIF2AKs activities (Krishna and Kumar, 2018) and PP1c mediated dephosphorylation. Importantly, and conversely to GADD34-deficient DCs, PERK deletion caused an increase in translation initiation and protein synthesis intensity as measured by SunRISE in both steady-state and LPS stimulated cDC2 (Fig. 6D). PERK is therefore the EIF2AK responsible for the majority of eIF2α phosphorylation in Flt3-L BMDCs DCs, and mirrors GADD34 activity to regulate active protein synthesis at steady state and during DC activation. These features presumably contribute to the acquisition of their immuno-stimulatory functions by DCs.

**Figure 6.**
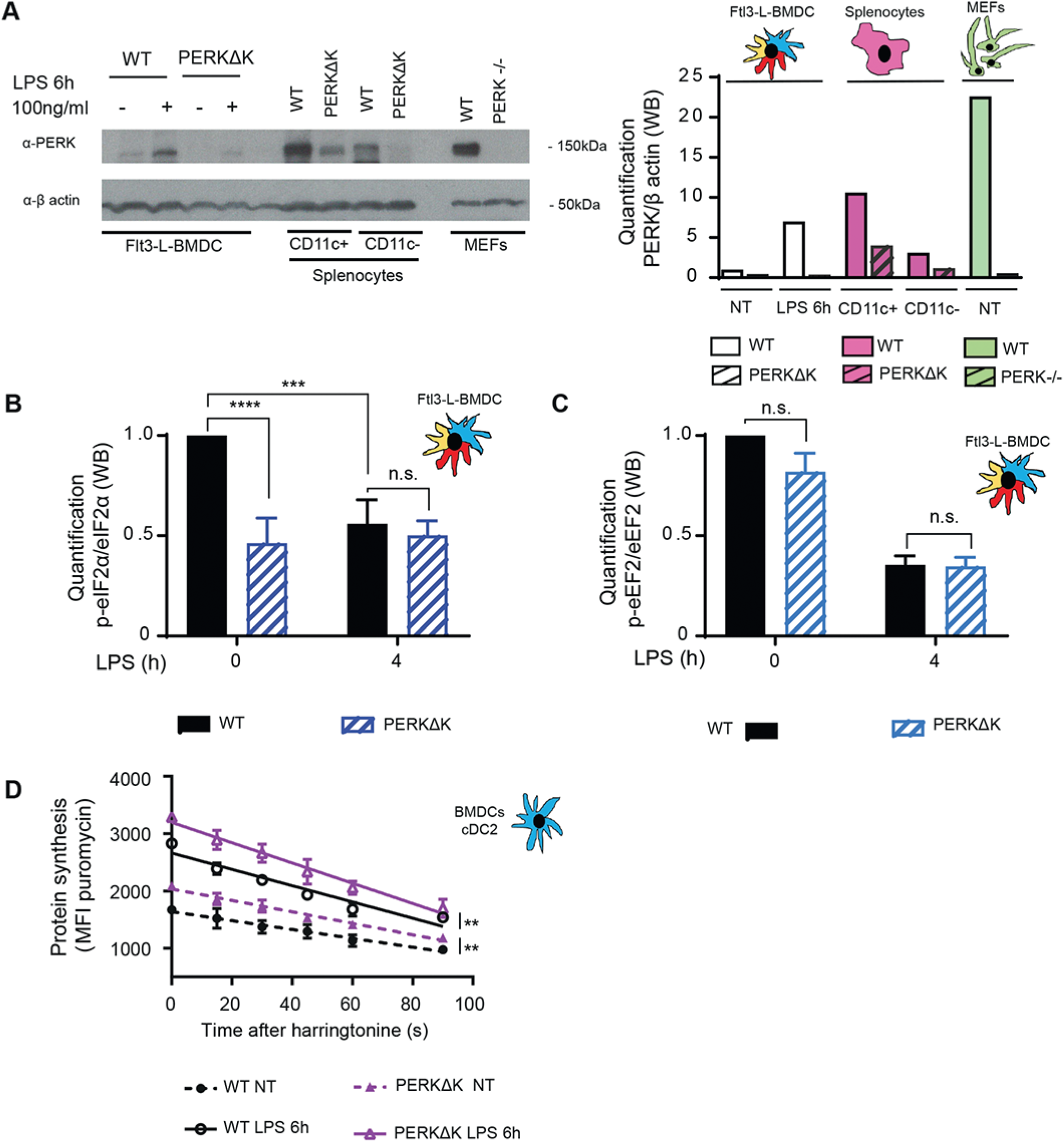
PERK is activated in steady-state dendritic cells. (A) Immunoblot detection of PERK and β-actin. Quantification is shown on the right. WT and PERKΔK Flt3-L BMDC treated or not with LPS (100ng/ml) for 6h were compared with CD11c+ and CD11c-fractions of splenocytes. WT and PERK-/- MEFs were used as control. (B-D) WT and Flt3-L PERKΔK BMDCs were stimulated or not with LPS (100ng/ml) for 4h. (B) Quantification of p-eIF2α/eIF2α ratio (immunoblot) in Flt3-L BMDCs. (C) Quantification of p-eEF2/eEF2 ratio (immunoblot) in Flt3-L BMDCs. Non-activated PERKΔK DCs display lower p-eIF2α levels than control cells. (D) The speed of translation elongation was measured using SunRISE in cDC2 after 4h of incubation with LPS. The total decay of puromycin MFI between WT and PERKΔK in steady-state and upon activation is showed. All data are representative of n=3 independent experiments. Data in (D) represent MFI +/- SD of 3 independent experiments. Unpaired Student’s t-test (** *p* < 0.01, *** *p* < 0.001 and **** *p* < 0.0001).

### DCs do not display chronic ISR features, but are resistant to ISR induction by subtilase cytotoxin

PERK is activated during DC development leading to intense eIF2α phosphorylation at steady-state, while these cells avoid translational arrest, by expressing GADD34 and eIF2B, among other potential compensatory biochemical mechanisms. We tested if an active UPR was induced during DC development, consequently inducing PERK dimerization and activation. First the splicing of XBP1 reflecting the activation of the IRE1α pathway (Walter and Ron, 2011) was found limited in the bulk of differentiating DCs (Sup Fig. S5A). In contrast to MEFs undergoing ER stress upon exposure to thapsigargin showed efficient splicing of XBP1. Similar observations were made for CHOP expression, which is one of the major transcription factors induced during the UPR (Han et al., 2013). Transcription factors ATF4 and ATF6 mRNA expression increased moderately during DC differentiation, their activation occurring mostly at the post-transcriptional level during stress (Walter and Ron, 2011). We next wondered if the relatively high PERK activity observed in developing DCs, could induce a chronic ISR in these cells. Recently, translational and transcriptional programs that allows adaptation to chronic ER stress has been described by Guan et al. (Guan et al., 2017). This chronic ISR (cISR) operates via PERK-dependent mechanisms, which allow simultaneous activation of stress-sensing and adaptive responses while allowing recovery of protein synthesis. Given the biochemical similarities with artificial cISR displayed by steady state DCs, we took advantage of available transcriptomic data (GSE9810, GSE2389) and of ATF4/CHOP-dependent gene (Han et al., 2013) to perform a Gene Set Enrichment Analysis (GSEA) and define the level of common gene expression found in DCs and potentially shared with a cISR. GSEA was followed by multiple testing correction (Subramanian et al., 2005) using the BubbleGUM software, which allows statistical assessment and visualization of changes in the expression of a pre-defined set of genes in different conditions (Spinelli et al., 2015). GSEA revealed no significant enrichment of ATF4- and CHOP-dependent genes expression in the DC transcriptome (FDR >0.25) (Sup Fig. S6A and S6B). Next, we compared cISR dependent transcription (Guan et al., 2017) with steady state splenocytes transcriptome, and found again no significant enrichment of cISR genes in the DC transcriptome compared to reference samples exposed to 1h or 16h of thapsigargin (Sup Fig. S6C and S6D) (FDR >0.25). DCs have therefore adapted to the consequences of high eIF2α phosphorylation to allow for translation of the DC transcriptome, without induction of chronic ISR. PERK- and eIF2α-P-mediated translational reprogramming during cISR appears to bypass cap-mediated translation and therefore to be largely eIF4F independent (Guan et al., 2017). We therefore used 4EGI-1, an inhibitor that interferes with eIF4F assembly (Moerke et al., 2007), and ROCA, an eIF4A inhibitor (Iwasaki et al., 2019), to treat WT and PERK-deficient DCs and confirm the dependency of their protein synthesis on eIF4F activity. Both small compounds had a profound inhibitory effect on DCs translational activity (80% of reduction), irrespective of their subsets or activation state (Sup Fig. S5B and S5C). This important level of inhibition indicates that DC mostly depend on eIF4F-dependent cap-mediated translation, thus again contrasting from cells undergoing chronic ISR (Guan et al., 2017). Despite high p-eIF2α levels, steady state DCs do not display a cISR-like response, ATF4-or CHOP-dependent gene transcription programs being unlikely part of DC development and function.

This lack of ATF4-dependent gene signatures in DCs when compared with other CD45+ cell types, lead us to wonder whether the high eIF2-P levels observed at steady state could interfere with DC capacity to respond to ER-stress. We thus exposed Flt3-L BMDCs to subtilase cytotoxin (SubAB), a bacterial AB5 toxin that promotes the release of the ER-chaperone BiP (HSPA5, Heat Shock Protein Family A (Hsp70) Member 5) and a PERK-dependent ISR (Paton et al., 2006). When WT and PERKΔK DCs were submitted to SubAB treatment a modest PERK-dependent induction of p-eIF2α was observed in WT cells (Fig. 7A), with limited consequences on translation (Fig. 7B). Conversely, thapsigargin treatment, arrested translation efficiently and triggered eIF2α phosphorylation potentially through a different EIF2AK than PERK, since high p-eIF2α levels were induced in PERKΔK cells (Fig. 7A and B). Little ATF4, could be detected in the cytosolic or nuclear fractions of control or toxin-treated DC, reflecting the modest induction of eIF2α phosphorylation (Fig. 7C and 7D), and confirming the limited impact of SubAB on DC translation and ISR induction. The efficacy of SubAB treatment was tested in MEFs, in which ATF4 was strongly induced by the toxin, and of course absent from control ATF4 -/- cell (Fig. 7C). DCs are therefore resistant to SubAB treatment and are unable to induce the ISR, despite the activation of other UPR branches, as demonstrated by augmented IRE1α-dependent splicing of the XBP1 mRNA (Fig. 7E). BiP mRNA levels were moderately augmented during DC differentiation. However, the similar expression ranges observed for DCs and MEFs (Fig. 7F) suggest that BiP transcriptional regulation is not involved in the DC resistance to SubAB. The relatively high PERK and GADD34 activity observed in steady-state DCs, together with high eIF2B expression, seems therefore to prevent the consequences of further activation of the ER kinase, leading to a minor increase in eIF2α phosphorylation and avoiding full ISR induction, translation arrest and associated ATF4 and CHOP synthesis.

**Figure 7.**
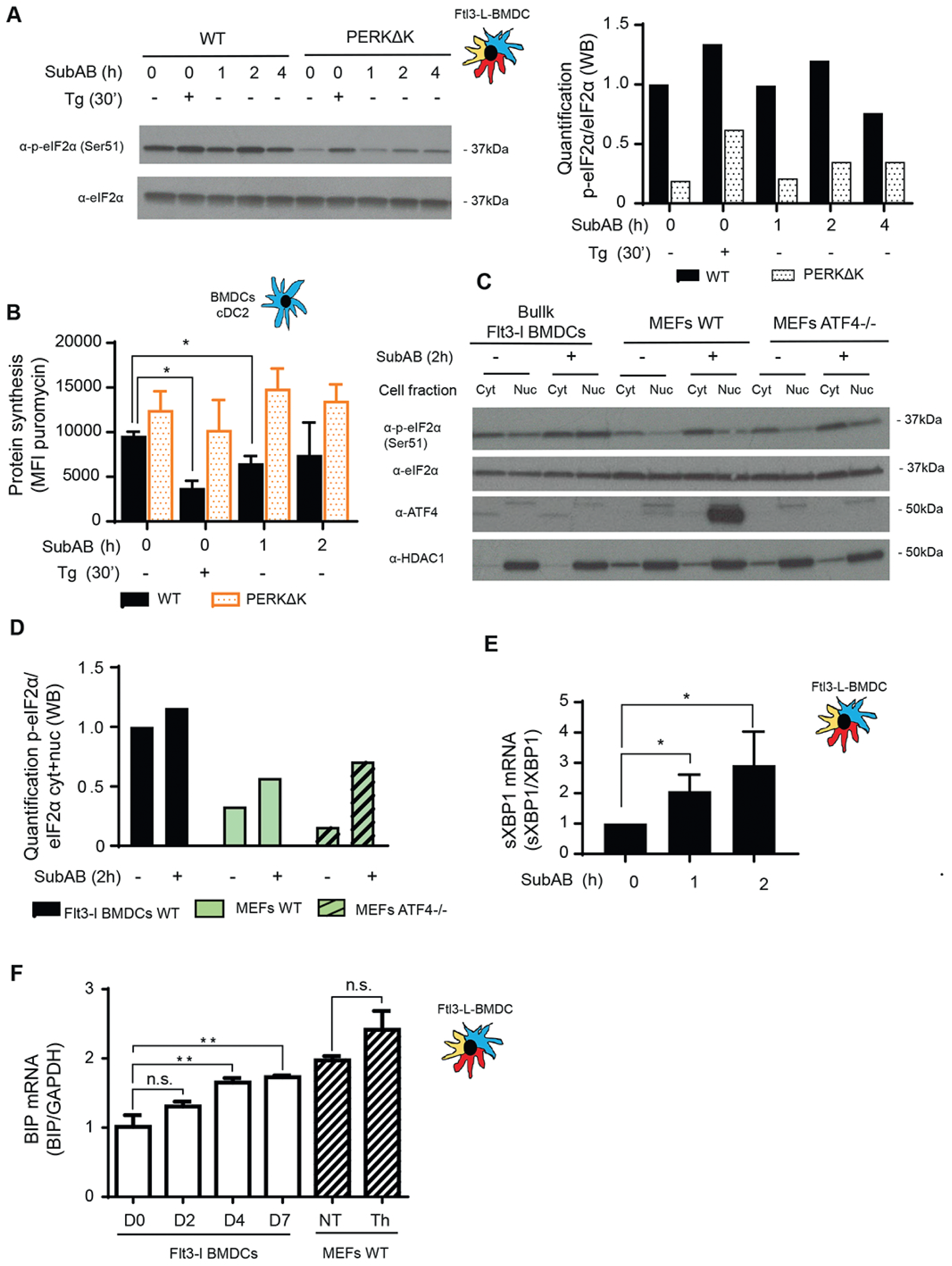
Flt3-L BMDCs are resistant to Subtilase cytotoxin-induced ISR. WT and PERKΔK Flt3-L BMDCs were stimulated with thapsigargin (200nM) and Subtilase cytotoxin (SubAB, 250ng/ml) for the indicated times. (A) Levels of p-eIF2α and total eIF2α detected by immunoblot (left) and quantified (right). (B) Levels of protein synthesis measured by flow using puromycilation detection in cDC2. Cells were incubated with puromycin 10 min before harvesting. The graph shows the total puromycin MFI levels. (C) WT Flt3-L BMDCs, WT and ATF4-/- MEFs were stimulated with Subtilase cytotoxin (SubAB, 250ng/ml) for 2h. Both cytoplasmic (cyt) and nuclear (nuc) fractions were analyzed. Levels of p-eIF2α, total eIF2α, ATF4 and HDAC1 (nuclear loading control) were revealed by immunoblot. p-eIF2α quantification is represented in (D). (E) WT Flt3-L BMDCs were stimulated with Subtilase cytotoxin (SubAB, 250ng/ml) for the indicated times. mRNA levels of spliced XBP1 were measured by RT-qPCR in bulk Flt3-L BMDCs. Raw data was normalized to total XBP1 mRNA expression. (F) Levels of BiP mRNA expression measured by RT-qPCR during Flt3-L BMDCs differentiation and in MEFs stimulated with thapsigargin for 30 min. Data represent MFI +/- SD of 3 independent experiments. Unpaired Student’s t-test (* *p* < 0.05, ** *p* < 0.01).

### Importance of the ISR for innate receptors signaling in DCs

In macrophages, p-eIF2α- and ATF4-dependent expression of HSPB8 is required for the assembly of signaling adaptors such as mitochondrial antiviral-signaling protein (MAVS), or TIR domain–containing adapter protein inducing interferon-α (TRIF), but not of Myeloid Differentiation Primary Response Gene 88 (MyD88). HSPB8 seems necessary for TRIF and MAVS to be incorporated into protein aggregates that constitute signalosomes for different innate immunity signaling pathways triggered by MAMPs (Abdel-Nour et al., 2019). Given the lack of ATF4 synthesis and the resistance of active DCs to mount an acute ISR, we investigated their capacity to produce pro-inflammatory cytokines and type-I IFN in a perturbed eIF2α-phosphorylation context. Importantly, we have previously shown that GADD34-deficient GM-CSF BM-derived and spleen DCs, have a reduced capacity to produce different cytokines, like IFN-β (Clavarino et al., 2012b, Perego et al., 2018), further suggesting that GADD34 activity and potentially eIF2α dephosphorylation is an important process for TLR signaling and cytokine production in DCs, although contrasting with the data obtained in macrophages (Abdel-Nour et al., 2019). LPS and the dsRNA synthetic mimic polyriboinosinic:polyribocytidylic acid (poly(I:C)), that is recognized by TRIF-dependent TLR3 and the MAVS-dependent cytosolic RNA helicases RIG-I and MDA5 (RLRs) (Fitzgerald and Kagan, 2020, Ablasser and Hur, 2020) were used to stimulate DCs (Fig. 8). Treatment with ISRIB, a pharmacological ISR inhibitor (Sidrauski et al., 2015), that prevents inhibition of eIF2B and promotes translation initiation (Fig. 8A), was included to prevent the induction of the ISR in activated DCs, and potentially interfere with cytokines expression, as described for poly(I:C)-stimulated macrophages (Abdel-Nour et al., 2019). ISRIB-treated DCs, however, did not show any relevant incapacitation in IFN-β nor IL-6 expression upon stimulation with either LPS or poly(I:C) (Fig. 8B and 8C). As expected, TLR4/Myd88 dependent LPS-activation of control bone marrow derived macrophages (BMM) was not impacted by ISRIB treatment (Abdel-Nour et al., 2019), while comparatively, poly I:C activation of these cells was too inefficient to obtain statistically reliable data (Fig. 8C). Thus, differently from macrophages, poly(I:C)-activated DCs do not require acute ISR induction, nor ATF4-dependent transcription to promote signalosomes assembly and transduce TRIF- or MAVS-dependent signaling to promote cytokines expression. Given the absence of impact of ISRIB on cytokine expression, we decided to analyze further the response to LPS of PERKΔK-DCs. This was prompted by the recent observation that PERK and IRE1α pharmacological inactivation affects Interleukin-1 beta (IL-1β) secretion in BMM. PERK was proposed to control the caspase-1-dependent proteolysis of pro-to mature-IL-1β, while IRE1α to keep mature IL-1β soluble to allow its secretion (Chiritoiu et al., 2019). LPS-activated WT and PERKΔK-DCs were co-stimulated with ATP to promote conversion of cytosolic pro-to mature IL-1β and its secretion (Fig. 8D). Surprisingly, we did not observe any impairment of IL-1β expression in PERKΔK-DCs, but rather an increase by 25% of both IL-1β mRNA transcription and mature IL-1β secretion compared to WT cells (Fig. 8D). This increased expression implies that PERK inactivation in DCs is not detrimental to IL-1β processing, but on the contrary, likely reflects the general protein synthesis augmentation displayed by the PERKΔK-DCs (Fig. 6D). The positive feedback loop created by augmented IL-1β secretion, could in turn increase *IL1B* mRNA transcription (Ceppi et al., 2009). Thus, conversely to macrophages, DCs do not rely on the ISR nor on PERK activity to produce cytokines in response to MAMPs, despite high p-eIF2α levels, that seems to prevent these cells to mount an acute stress response.

**Figure 8.**
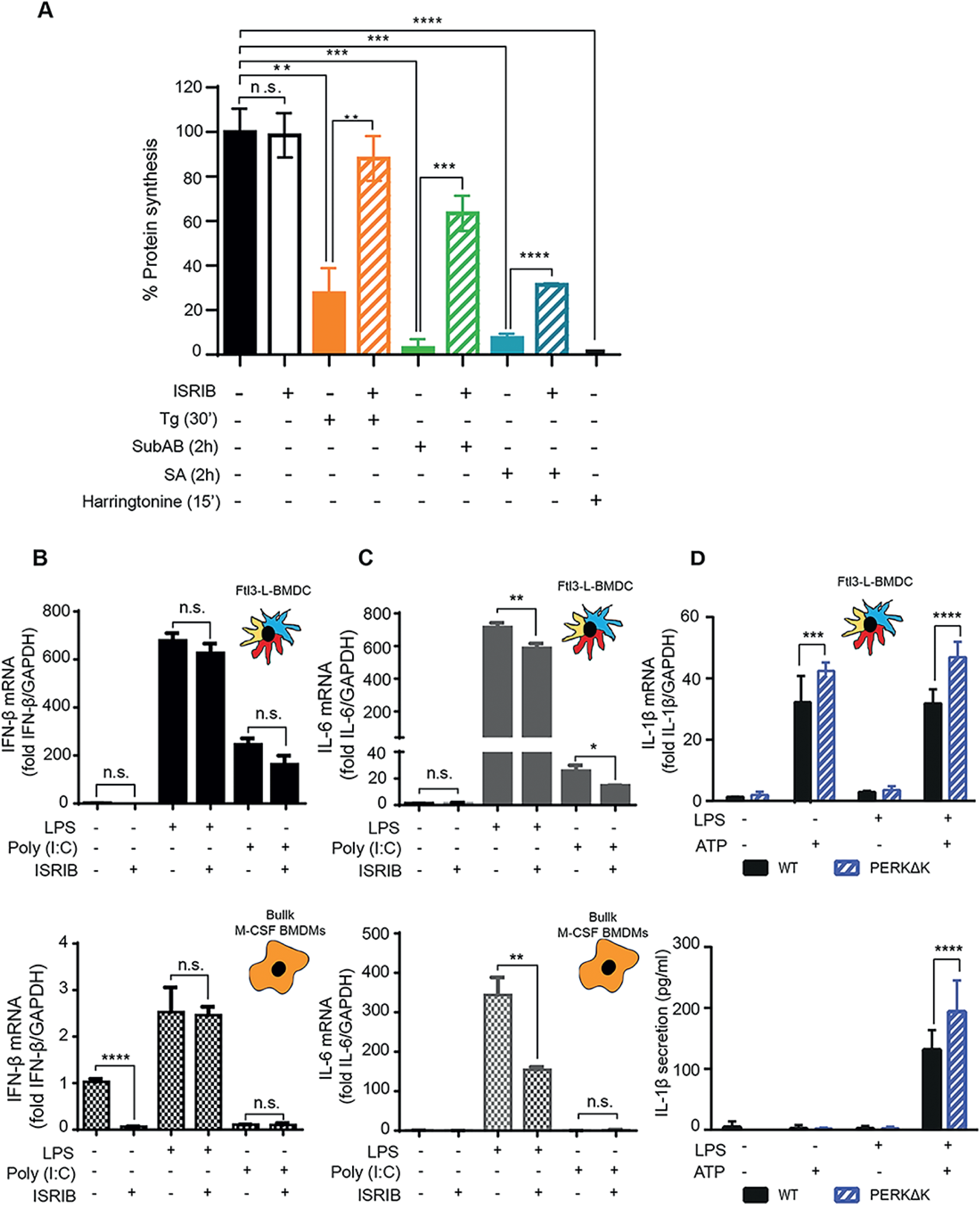
ISRIB does not inhibit cytokines expression in MAMPs activated Flt3-L BMDCs. (A) Protein synthesis was measured by puromycilation and flow in MEFs treated with different ISR inducing and ISRIB drugs for indicated times. ISRIB treatment was efficient to prevent protein synthesis inhibition upon eIF2α phosphorylation induction by thapsigargin (Tg), Subtilase cytotoxin (SubAB) and Sodium Arsenite (SA). (B) IFN-β mRNA expression measured by RT-qPCR in Flt3-L BMDCs and BMM activated with LPS or Poly (I:C) and treated or not with ISRIB for 4 hours. (C) IL-6 mRNA expression measured by RT-qPCR in Flt3-L BMDCs and BMM activated with LPS or Poly (I:C) and treated or not with ISRIB. (D) IL-1β mRNA expression measured by RT-qPCR (left) and secretion by ELISA (right) in WT and PERKΔK Flt3-L BMDCs stimulated with LPS during 4h and with ATP for the last 30 minutes of treatment. Data represent MFI +/- SD of 3 independent experiments. Unpaired Student’s t-test (** *p* < 0.01, *** *p* < 0.001 and **** *p* < 0.0001).

### PERK and actin polymerization coordinates p-eIF2α levels and migration in DCs

Recently, the importance of globular actin in the formation of a tripartite holophosphatase complex assembled with GADD34 and PP1c to dephosphorylate eIF2α was revealed (Crespillo-Casado et al., 2017, Chambers et al., 2015, Crespillo-Casado et al., 2018). Given the unusual regulation of eIF2α phosphorylation in DCs, we wondered whether actin organization could also impact this pathway in a different setting than artificial ER-stress induction. Actin depolymerizing and polymerizing drugs, respectively Latrunculin A (Lat A) and Jasplakinoloide (Jaspk) had opposite effects on eIF2α phosphorylation in Flt3-L BMDCs (Fig. 9A). Globular actin accumulation induced by Lat A was strongly correlated with eIF2α dephosphorylation (Fig. 9A and 9B), while actin polymerization induced by Jaspk resulted in a massive increase of eIF2α phosphorylation, together with a reduction in protein synthesis (Fig. 9A and 9C), similarly to what was observed in GADD34-deficient DCs. We next tested the impact of the two drugs on WT and PERKΔK Flt3-L BMDCs activated or not by LPS (Fig. 9D). LPS activation or Lat A treatment resulted in the same levels of eIF2α dephosphorylation and had a synergistic effect when used together (Fig. 9D). Jaspk dominated the LPS effect and strongly increased p-eIF2α levels in all conditions tested. PERK inactivation decreased the levels of p-eIF2α, but had no obvious consequences on the efficacy of the drugs, since both induced similar responses in WT DCs. We next tested the impact of actin remodeling and translation regulation on the acquisition by DC of their immune-stimulatory phenotypes. Surface MHC II and co-stimulatory molecule CD86 expression were up-regulated by LPS stimulation but remained unaffected by Jaspk treatment (Fig. 9E). Similarly, transcription levels of key cytokines like IL-6 and IFN-β were insensitive to this actin polymerizing drug (Fig. 9F & 9G). However, while IL-6 secretion remained identical, IFN-β levels were found reduced by Jaspk treatment, suggesting that IL-6 mRNA translation occurs normally despite high eIF2α-phosphorylation conditions. Interestingly, the IL-6 gene was described to bear an upstream uORF-dependent translational regulation, that could allow IL-6 mRNA translation during Jaspk treatment (Sanchez et al., 2019). Taken together, these observations suggest that extensive actin polymerization in DCs increases strongly eIF2α phosphorylation, affecting protein synthesis and ultimately controlling translationally specific cytokines expression, similarly to what has been observed with Cdc42 or Wiskott-Aldrich syndrome protein (WASp) mutants (Pulecio et al., 2010, Prete et al., 2013). These results further suggest that actin dynamics and its effect on eIF2α phosphorylation could be a key regulating factors for specific mRNAs homeostasis and translation, like IFNs, that are generally secreted in a polarized fashion or needs to be coordinated with cell migration (Pulecio et al., 2010, Prete et al., 2013). Finally, given the interplay between eIF2α phosphorylation and actin polymerization, we wondered whether PERK-deficient cells could display some migratory deficits. We used microfabricated channels, that mimic the confined geometry of the interstitial space in tissues (Heuze et al., 2013, Bretou et al., 2017), to find that PERK-deficient cells were not able to increase their migration speed in response to LPS (Fig. 9G), confirming the link between eIF2α phosphorylation and actin dynamics. PERK activity is therefore necessary for DCs to acquire normal immune-stimulatory and migratory activities, presumably by coordinating protein synthesis and translation specific mRNAs with actin polymerization.

**Figure 9.**
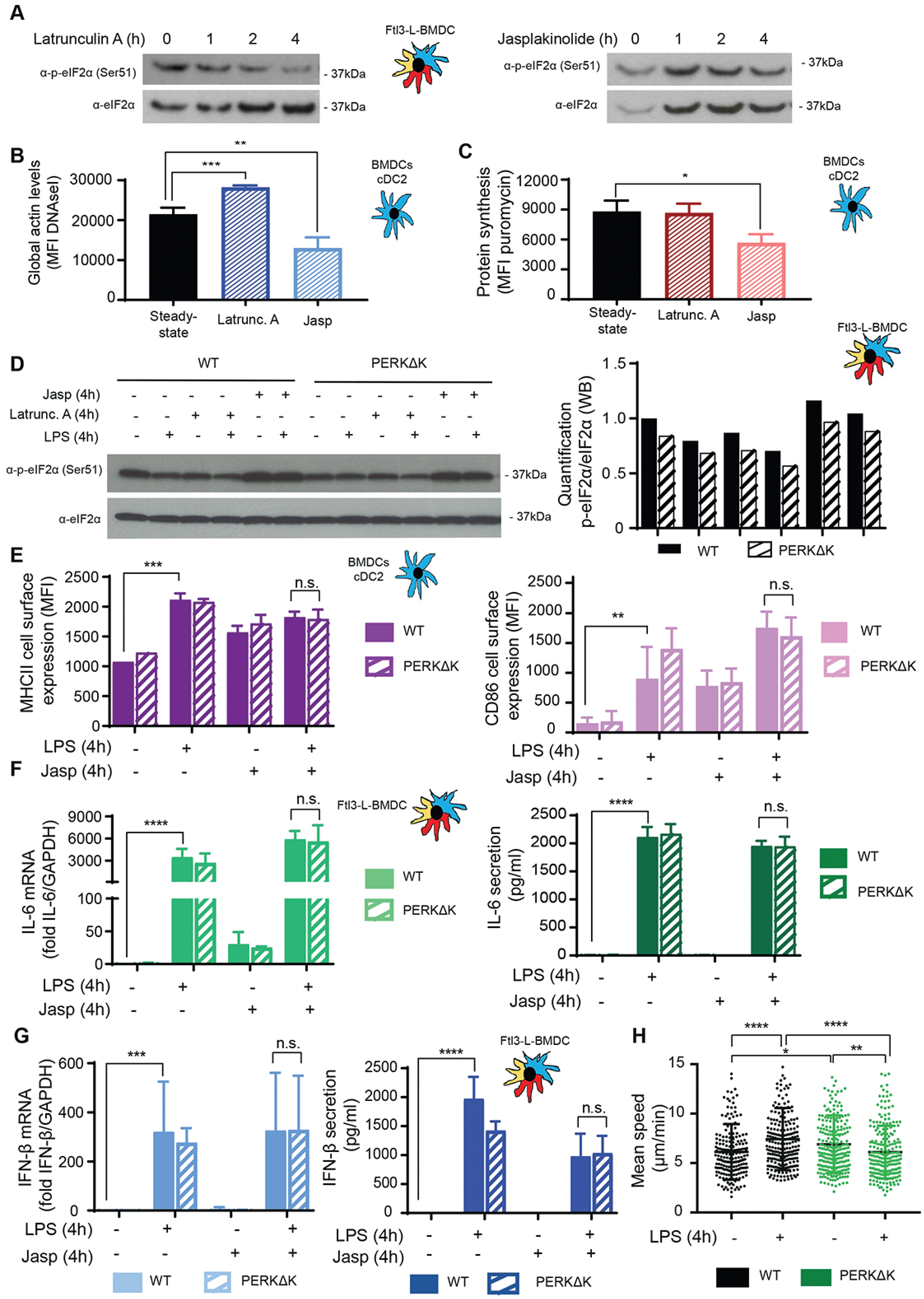
eIF2α phosphorylation levels are regulated by G-actin availability. WT Flt3-L BMDCs were treated with Latrunculin A (50nM) (Latrunc A) and Jasplakinolide (1µM) (Jasp) for the indicated times. (A) Levels of p-eIF2α and total eIF2α detected by immunoblot in Flt3-L BMDCs. (B) Globular actin levels were measured by flow cytometry intracellular stainning using a fluochrome coupled DNAse I protein in cDC2 population from Flt3-L BMDCs. The graph shows total MFI levels. (C) Levels of protein synthesis in cDC2 were measured by puromycilation and flow intracellular staining. Cells were incubated with puromycin 10 min before harvesting. The graph shows total puromycin MFI levels. WT and PERKΔK Flt3-L BMDCs were treated with LPS (100ng/ml), Latrunc A and Jasp for 4h. (D) Levels of p-eIF2α and total eIF2α monitored by immunoblot in Flt3-L BMDCs. Quantification is shown on the right. (E) Upregulation of surface MHC II and CD86 levels was measured by flow in cDC2. The graphs represent total MFIs. (F-G) mRNA levels and secretion of IL-6 (F) and IFN-β (G) were measured by RT-qPCR and by ELISA from Flt3-L BMDCs cultures. Raw data was normalized to the housekeeping gene (GAPDH). H) WT and PERKΔK GM-CSF BMDCs were treated with LPS (100ng/ml) during 30 min previous to 16h of migration. The graph represents instantaneous mean velocities of migration in 4-by 5µm fibronectin-coated microchannels of at least 100 cells per condition. All data are representative of n=3 independent experiments. Data are mean ± SD (n = 3). Unpaired Student’s t-test (* *p* < 0.05, ** *p* < 0.01, *** *p* < 0.001 and **** *p* < 0.0001).

## Discussion

We have previously proposed the existence of a strong causative link between cell activation by TLR ligands and eIF2α phosphorylation, notably by virtue of strong GADD34 expression in most transcriptomics studies performed on PAMPs-activated DCs (Claudio et al., 2013, Clavarino et al., 2012b, Reverendo et al., 2018). Our present work suggests that steady state DCs activate PERK-mediated eIF2α phosphorylation to acquire their functional properties during differentiation (Sup Fig. 7), but distinctly from known ISR programs, normally induced upon acute or chronic ER stress (Han et al., 2013, Guan et al., 2017).

To our knowledge, the level of p-eIF2α observed in primary DC both *in vivo* and *in vitro* are unique in their amplitude. Although, as judged comparatively from experiments performed with artificial induction of the different EIF2KAs, such p-eIF2α levels should be inhibitory for global protein synthesis (Dalet et al., 2017). DCs have acquired biochemical resistance, like high expression of eIF2B, to compensate for the consequences of this developmental PERK activation and to undergo high eIF2α phosphorylation while maintaining normal proteostasis. The role of ATF4 in controlling *Ppp1r15a/GADD34* mRNA expression and eIF2α dephosphorylation to restore protein synthesis during stress has been extensively studied (Novoa et al., 2001), but the active translation observed in DCs is likely not permitting ATF4 synthesis and consequently the activation of a bona fide ISR despite high eIF2α phosphorylation. GADD34 is nevertheless functional in non-activated DCs, in which IKKϵ/TBK1 activity and potentially the transcription factor IRF3 are required for its mRNA transcription. This dependency of *Ppp1r15a/GADD34* transcription on IKKϵ/TBK1 confirms that the *PPP1R15a* gene belongs to a group of genes directly induced by TLR or RLR signaling, as previously suggested by ChIP-seq and transcriptomics analysis of viral or poly (I:C)-stimulated cells (Freaney et al., 2013, Lazear et al., 2013), Dalet et al., 2017). GADD34 protein expression is undetectable in DCs, without prior treatment with proteasome inhibitors (Clavarino et al., 2012b), which is presumably indicative of an extremely tight equilibrium between its translation and active degradation. GADD34 mRNA translation, like that of ATF4, is controlled through 5’ upstream ORFs regulation, which are normally bypassed upon global translation arrest to favor the synthesis of these specific ISR molecules (Palam et al., 2011). Interestingly, GADD34 mRNA has been recently shown to be also actively translated in unstressed MEFs, albeit at much lower levels than upon ER stress (Reid et al., 2016). Thus, in steady state DCs, GADD34 synthesis likely occurs in a high eIF2α-phosphorylation context, despite relatively normal level of translation, while that of ATF4 does not. The difference in the 5’ upstream ORFs organization of the mRNAs coding for these two molecules (Palam et al., 2011, Andreev et al., 2015), could explain this difference, and how GADD34/PP1c activity contributes to the maintenance of protein synthesis activity by counteracting PERK in steady state DCs.

Importantly, the PERK/eIF2α/GADD34 molecular trio sets the physiological range for potential protein synthesis initiation available in the different DC activation stage, however the upward progression triggered by LPS stimulation, from one level of protein synthesis to the next, does not depend on this biochemical axis and is likely regulated by other protein synthesis regulation pathways, like the mTORC1 or Casein Kinase 2 pathways (Lelouard et al., 2007, Reverendo et al., 2019) (Sup Fig. 7). Other mechanisms that contribute to escape PERK-mediated eIF2α phosphorylation, including the eIF2B-independent and eIF3-dependent pathway recently described to rescue translation during chronic ER stress (Guan et al., 2017), do not seem to be used in the DC context. These DC specific mechanisms prevent the induction of the ISR by the AB5 subtilase cytotoxin, which targets the ER chaperone BiP. Independently of demonstrating that induction of acute eIF2α phosphorylation by thapsigargin is not solely dependent on PERK activation, our observations suggest that DCs could escape EIF2KA-dependent translation arrest during exposure to different metabolic insults relevant to the immune context. These situations can include viral infection (Clavarino et al., 2012b), exposure to high levels of fatty acids during pathogenesis, oxidative stress during inflammation or amino acids starvation, mediated by amino acid degrading enzymes, like Arginase 1 or IDO, which are induced during infection or cancer development (Claudio et al., 2013, Munn et al., 2004). Importantly we could also show, that in contrast to macrophages, PERK and ISR induction are not necessary for DCs to drive the transcription of pro-inflammatory cytokines in response to TRIF or MAVs dependent-signaling (Abdel-Nour et al., 2019) nor the secretion of IL-1β (Chiritoiu et al., 2019).

PERK activation is therefore required to regulate mRNA translation during DC development and potentially also GADD34 synthesis, which not only provides a negative feed-back, but also is required for normal DC activation and cytokines expression (Clavarino et al., 2012b, Perego et al., 2018). Although a role for the ISR has been suggested to favor the survival of tissue associated DCs (Tavernier et al., 2017), we could not detect any particular phenotype impairing the development of DCs in the spleen of PERK-deficient animals. A close examination of the functional capacity of DC *in vitro* showed that in addition of a modified production of several cytokines, activated PERK-deficient DCs show an impaired migratory capacity. This finding echoes with the existence of a cross-talk between actin skeleton organization and protein synthesis regulation. We confirmed that globular actin synergizes with the PP1c to dephosphorylate p-eIF2α, suggesting that the PERK/GADD34 pathway could play an important role in regulating globally or locally translation in response to actin dynamics and possibly coordinating migration. DCs represent a model of choice for studying this possibility, given their developmental regulation of eIF2α phosphorylation and their requirement for actin dependent-phagocytosis and migration to perform their immune-stimulatory function. The activation of PERK/GADD34 pathway in steady state DC also underlines the importance of these molecules in homeostatic condition, independently of obvious acute ER stress, for the acquisition of specialized function. Our findings open therefore new pharmacological perspectives for therapeutic immune intervention by targeting PERK, GADD34 or eIF2α phosphorylation.

## Acknowledgements

We thank all CIML cytometry and Imaging core facilities for expert assistance. The laboratory is supported by grants from La Fondation ARC. The laboratory is “Equipe de la Fondation de la Recherche Médicale” (FRM) sponsored by the grant DEQ20140329536. The project was also supported by grants from l’Agence Nationale de la Recherche (ANR), « ANR-FCT 12-ISV3-0002-01» and « INFORM Labex ANR-11-LABEX-0054 », «DCBIOL Labex ANR-11-LABEX-0043 » and ANR-10-IDEX-0001-02 PSL* and A*MIDEX project ANR-11-IDEX-0001-02 funded by the “Investissements d’Avenir” French government program. The research was also supported by the Ilídio Pinho foundation and FCT - Fundação para a Ciência e a Tecnologia - and Programa Operacional Competitividade e Internacionalização - Compete2020 (FEDER) – references PTDC/IMI-IMU/3615/2014 and POCI-01-0145-FEDER-016768. We thank Lionel Spinelli and Thien-Phong Vu-Manh at CIML for bioinformatics and statistics support. We acknowledge financial support from n° ANR-10-INBS-04-01 France Bio Imaging and the ImagImm CIML imaging core facility. The authors declare to have no competing interest.

## AUTHORS’ CONTRIBUTION

A.M., J.G., D.B, S.C., D.S. V.C., R.J.A., L.C., C.R.R. performed research. A.P and J.P provided key reagents. A.M, A-M.L-D., E.G. and P.P. designed research and analyzed data. A.M, J.G., and P.P. wrote the paper.

## Figure legends

**Supplementary Figure 1.**
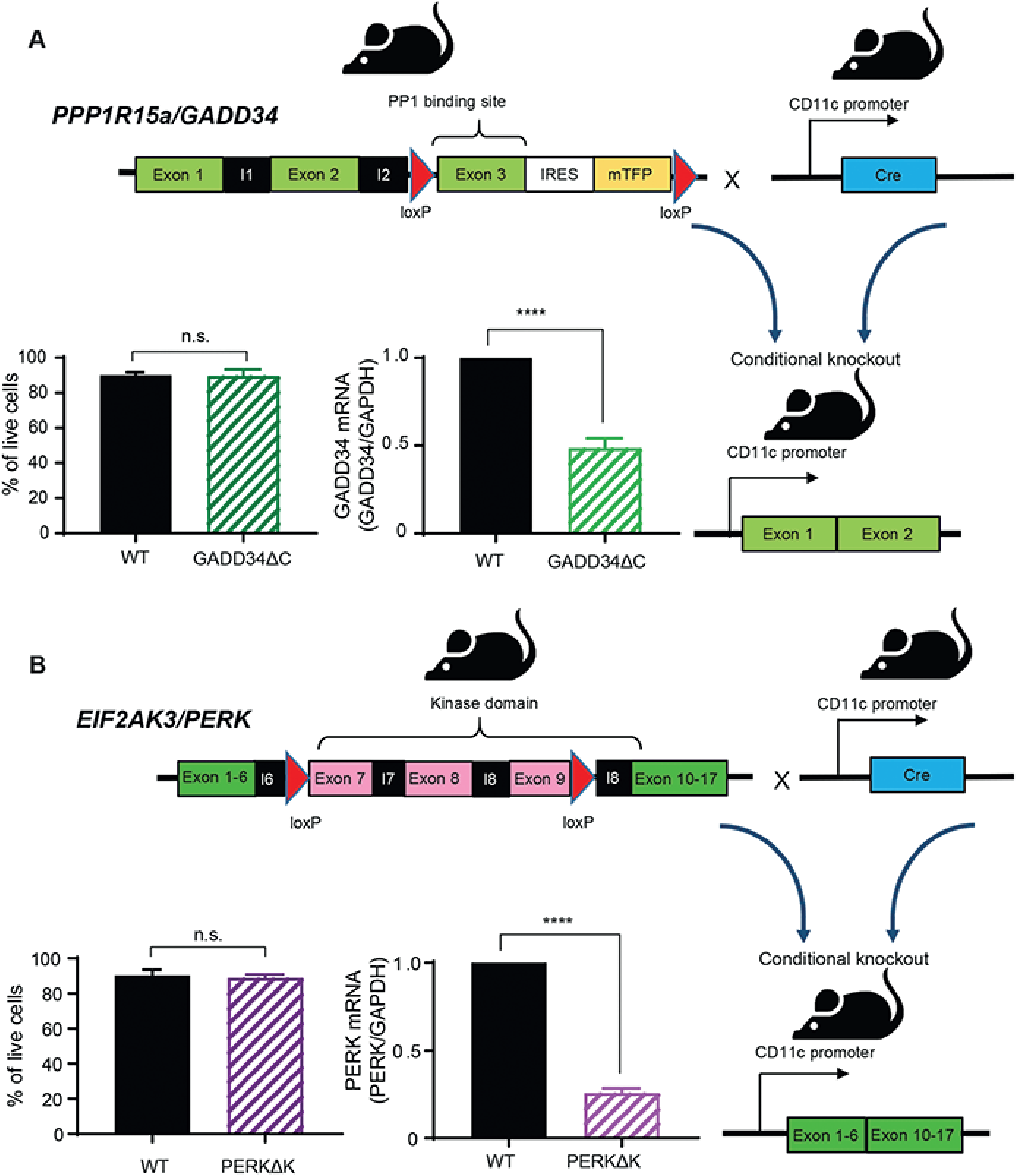
GADD34^loxp/loxp^ CD11c Cre+ mice and PERK^loxp/loxp^ CD11c Cre+ mice. (A) The *PPP1R15a* gene (GADD34) is constituted by three exons, and the third 3 contains the PP1c binding domain. GADD34ΔC^loxp/loxp^ CD11c Cre+ mice (GADD34ΔC) was generated using a combination of Cre/loxP and FRT/FLP systems. GADD34ΔC^loxp/loxp^ was developed by introducing a neomycin resistance cassette flanked by two loxP sequences in the third exon and a third loxP site in the end of the second exon. Mice were bred to Itgax-Cre+ mice (Caton et al., 2007) mice to generate GADD34ΔC in which the GADD34 gene is lacks the PP1 binding domain in cells expressing CD11c+. This excision event was confirmed by PCR in Flt3-L BMDCs populations from WT and GADD34ΔC mice. GADD34ΔC mice are viable and no difference was observed in the differentiation or survival of Flt3-L BMDCs *in vitro*. (B) The *EIF2KA3* gene (PERK) is constituted of 17 exons. Exons 7, 8 and 9 codes for part of the ER lumenal activation domain, the transmembrane domain and part of the cytoplasmic catalytic domain. Deletion of exons 7, 8 and 9, inactivates PERK by altering its ER localization and kinase domain. PERK^loxp/loxp^ mouse was generated by introducing a neomycin resistance cassette flanked by two loxP sequences in the end of the ninth exon and a third loxP site in the end of the sixth exon (Zhang et al., 2006). PERK^loxp/loxp^ mice were bred to Itgax-Cre+ mice to generate PERK^loxp/loxp^ CD11c Cre+ mice (PERKΔK). Excision of PERK floxed alleles was confirmed by PCR in Flt3-L BMDCs from WT and PERKΔK. PERKΔK mice are viable and no difference was observed in the survival of Flt3-L BMDCs.

**Supplementary figure 2.**
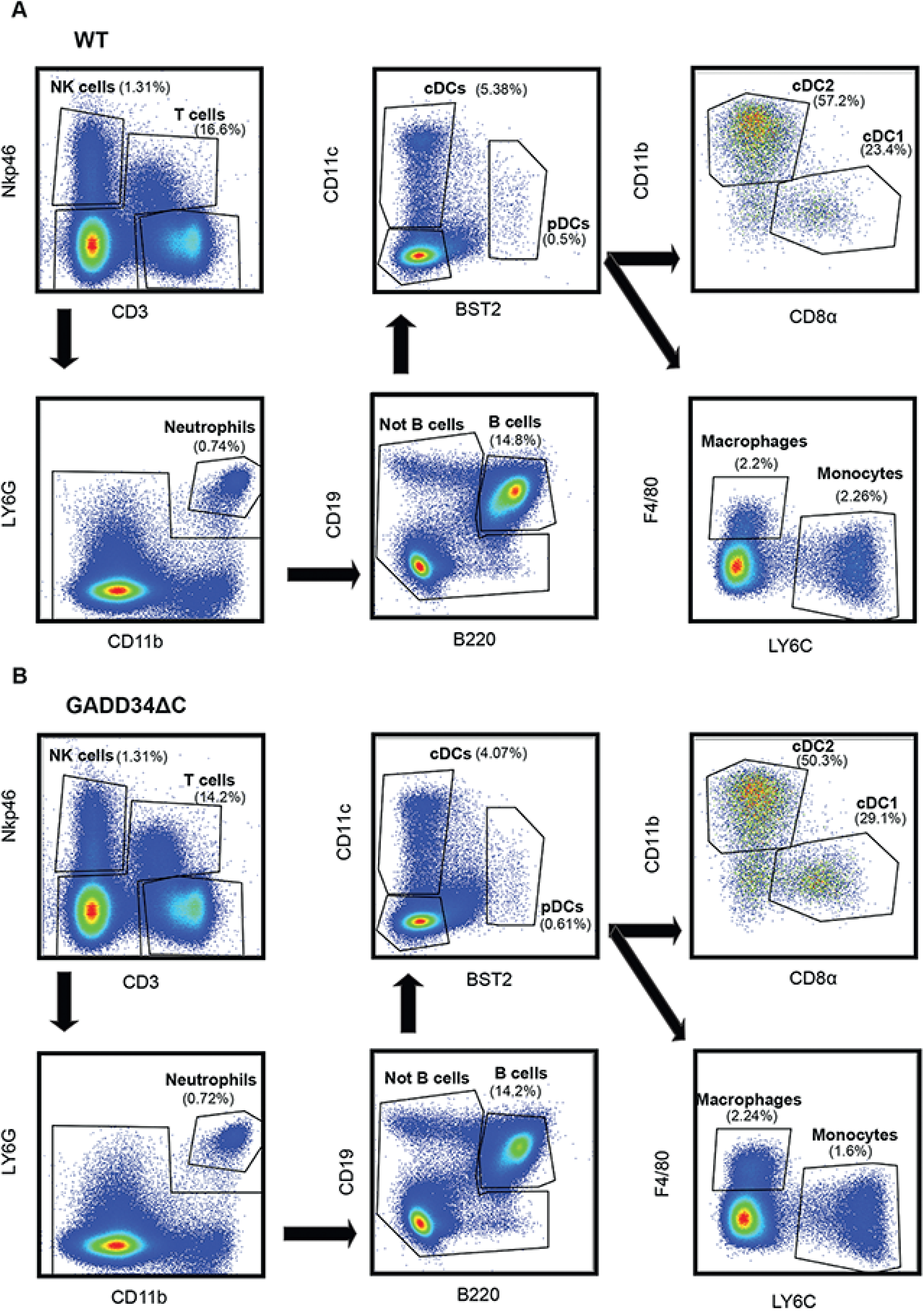
GADD34ΔC mouse spleen populations. Splenocytes from WT (A) and GADD34ΔC (B) mice were analyzed by flow cytometry. Splenocytes were identified based on Nkp46+ (NK cell) and CD3+ (T cells). We gated the double negatives in LY6G and CD11b, selecting the double-positives as neutrophils. The rest of the population was divided in B cells (B220 +) and not B cells (B220-). The remaining cells were classified as DCs (CD11c+ MHCII+), macrophages (CD11c-F4/80+) and monocytes (F4/80-LY6C+).

**Supplementary figure 3.**
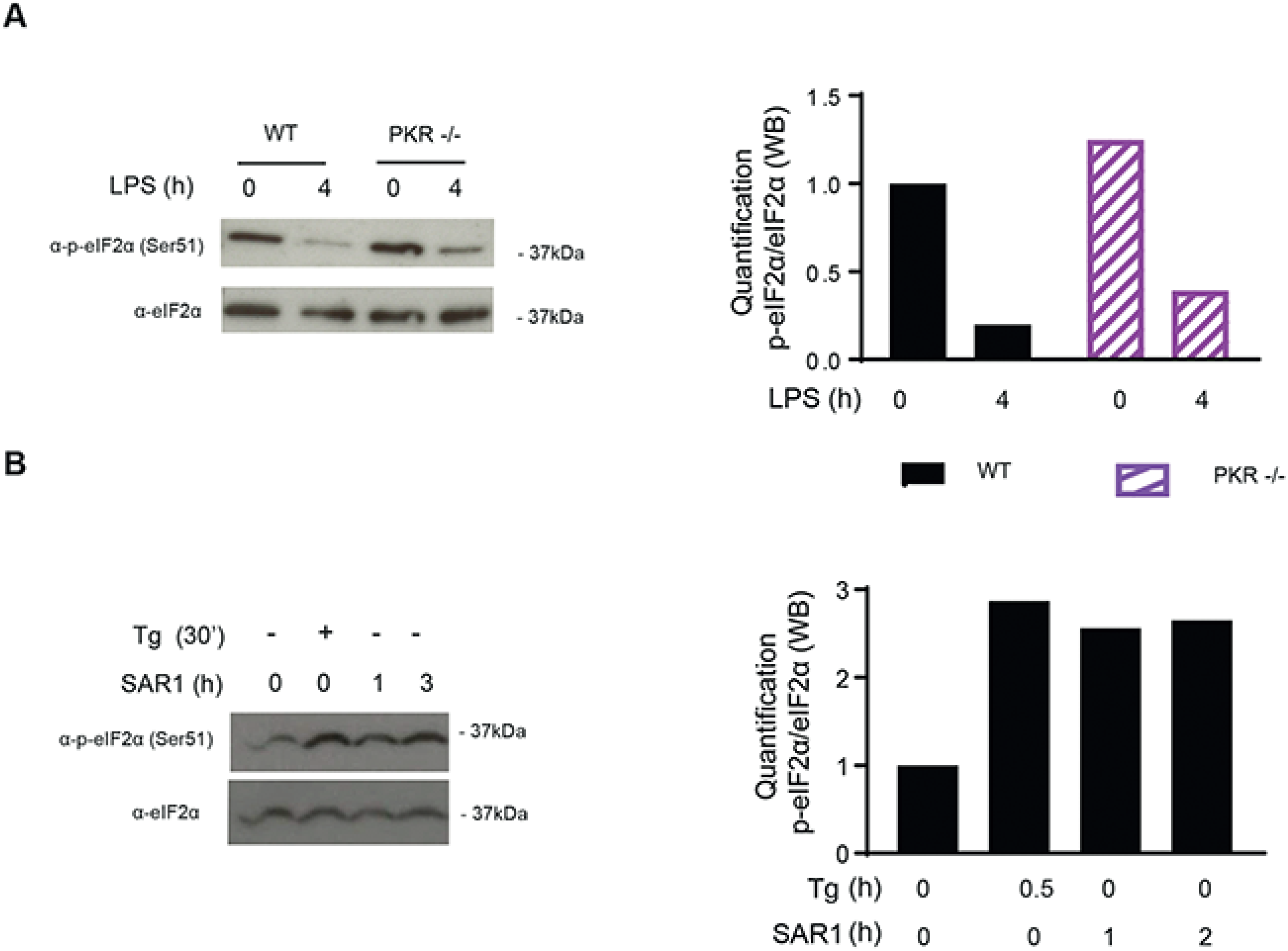
Inhibition of PKR and GCN2 does not alter steady-state p-eIF2α levels. (A) WT and PKR-/- Flt3-L BMDCs were stimulated with LPS (100ng/ml) during 4 hours. Levels of p-eIF2α and total eIF2α are shown by immunoblot and the respective quantification is represented in the graph on the right. (B) WT Flt3-L BMDCs were stimulated with SAR1, a specific inhibitor of PKR and GCN2 kinases (1µM) and thapsigargin (200nM) for the indicated times. Levels of p-eIF2α and total eIF2α are shown by immunoblot and the respective quantification is represented in the graph on the right.

**Supplementary figure 4.**
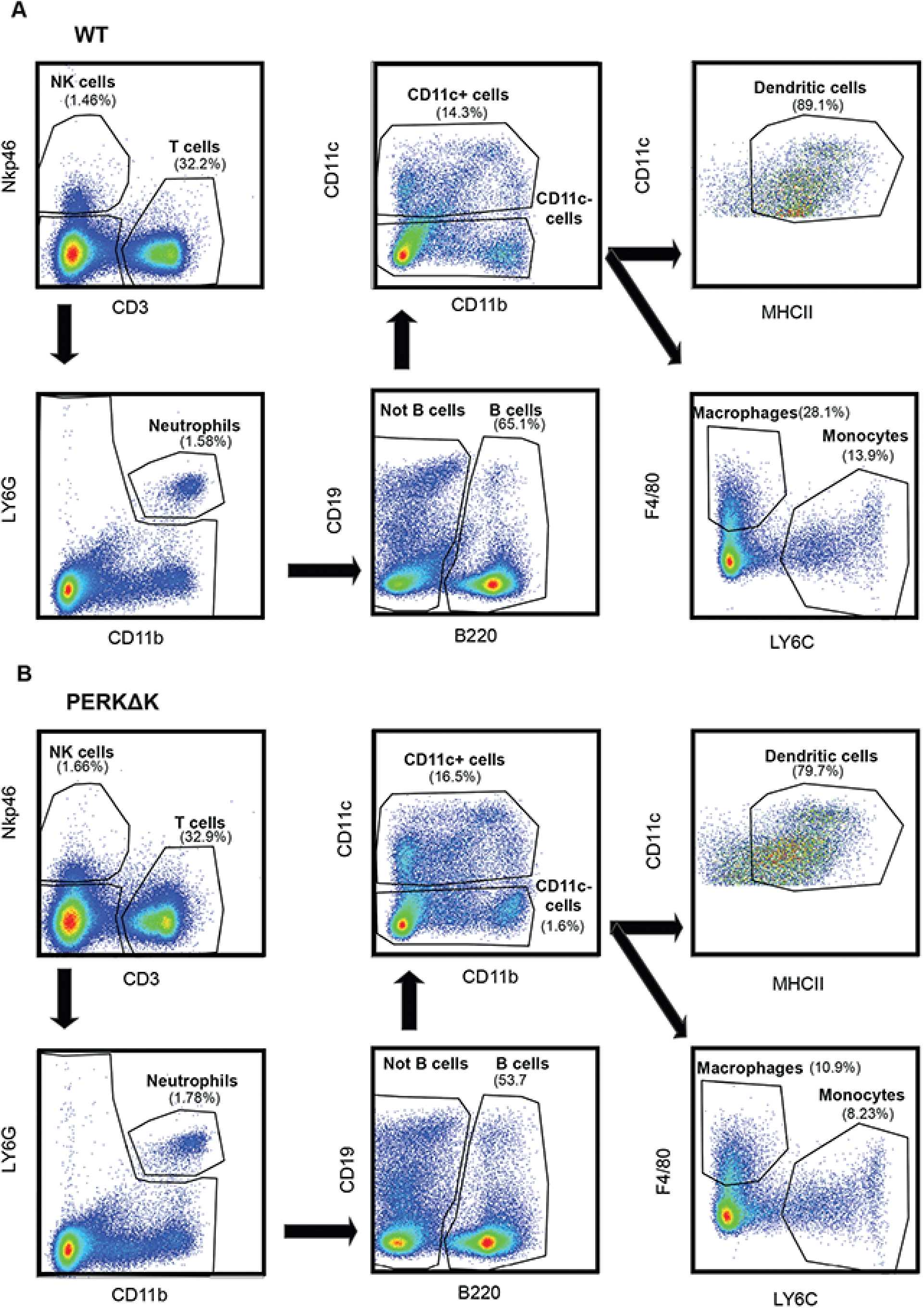
PERKΔK mouse spleen populations. Splenocytes from WT (A) and PERKΔK (B) mice were analyzed by flow cytometry. Splenocytes were identified based on Nkp46+ (NK cell) and CD3+ (T cells). We gated the double negatives in LY6G and CD11b, selecting the double-positives as neutrophils. The rest of the population was divided in B cells (B220 +) and not B cells (B220-). The remaining cells were classified as DCs (CD11c+ MHCII+), macrophages (CD11c-F4/80+) and monocytes (F4/80-LY6C+).

**Supplementary figure 5.**
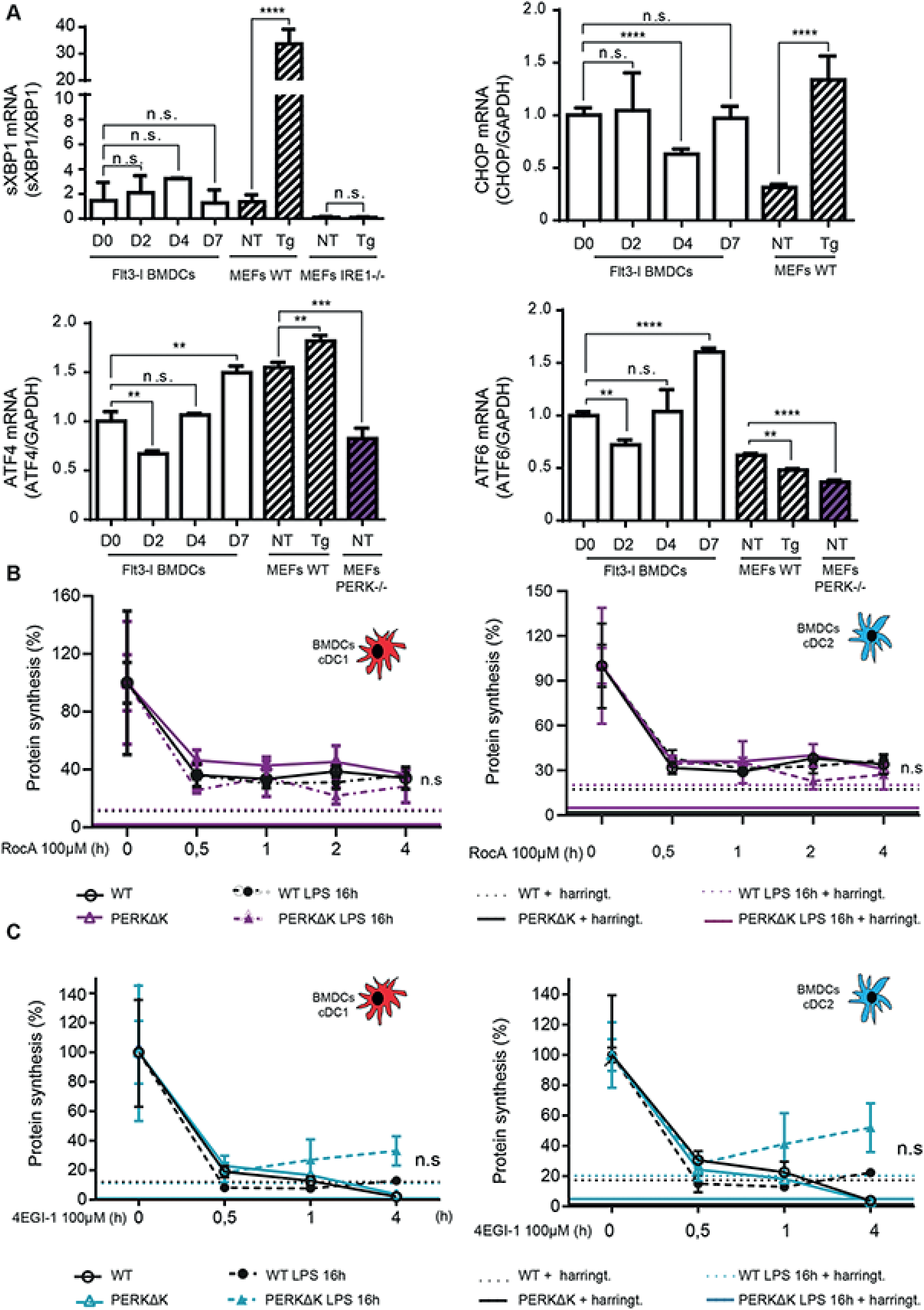
Steady state DCs display eIF4F-dependence for their protein synthesis. (A) Monitoring of XBP1s mRNA splicing, ATF6, ATF4 and CHOP mRNA expression by RT-qPCR during Flt3-L BMDCs differentiation and in MEFs. (B and C) Protein synthesis activity upon ROC1 (B) or 4EGI treatment of WT (left) and PERKΔK (right) Flt3-L BMDCs during time. DC do not display a chronic ISR, nor become eIF4G-independent for their translation, as suggested for long term exposure to thasigargin (Guan et al., 2017).

**Supplementary figure 6.**
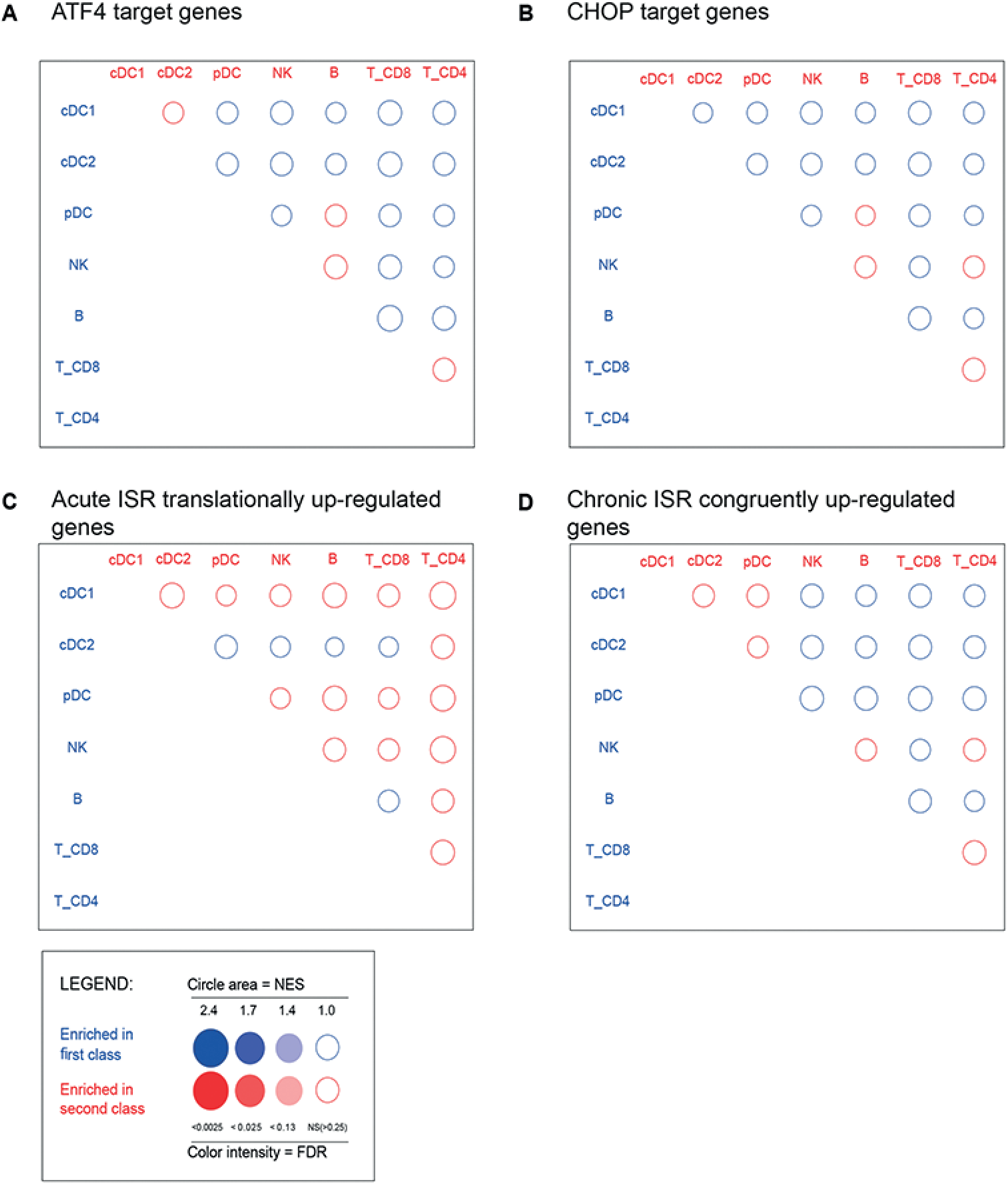
Gene set enrichment analyses (GSEA) of cISR and ATF4 regulated gene expression in DCs. No significant ISR-related gene signature can be observed in dendritic cells. GSEA using BubbleGUM (see methods) performed on: (A) ATF4-dependent genes. (B) CHOP-dependent genes. (C) Genes translationally up-regulated in macrophage after 1h of thapsigargin treatment. (D) Genes congruently up-regulated after 16h of thapsigargin treatment.

**Supplementary figure 7.**
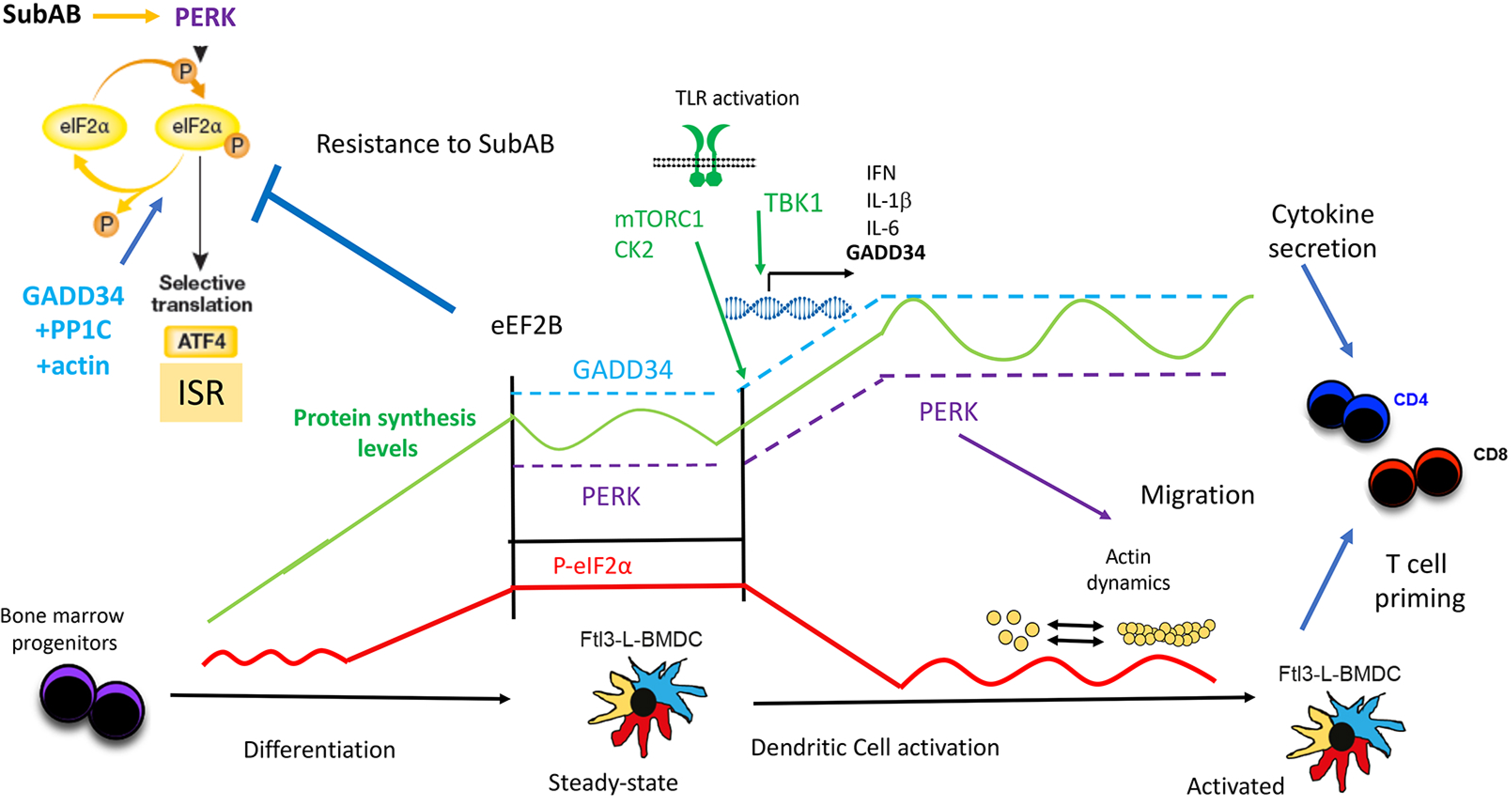
Graphical abstract. In most cells SubAB drives PERK to phosphorylate eIF2α, a dominant negative inhibitor of the GEF eIF2B, that inhibits global protein synthesis while promoting ATF4 synthesis, as well as, the Integrated Stress Response (ISR). The phosphatase-1 co-factor GADD34/PPP1R15a, in complex with PP1c and globular actin, promotes the dephosphorylation of eIF2α. During DC differentiation, PERK is activated and majorly responsible for eIF2α phosphorylation (red). Primary dendritic cells display therefore extremely high p-eIF2α levels comparatively to other immune cells. PERK and GADD34 control protein synthesis levels in steady state DCs (green), which are both increased during differentiation and upon TLR activation. PERK (purple) and GADD34 (cyan) maintain protein synthesis (green) in a window of activity relative to a set baseline, which changes during DC differentiation and activation. The intensity of protein synthesis baseline is controlled by mTORC1 and Casein Kinase 2 (CK2) signaling cascades, which are both activated upon LPS sensing, along with TBK1, which promotes the transcription of GADD34, Interferons and different cytokines genes (Lelouard et al. 2007, Reverendo et al, 2019). GADD34 and high eIF2B synthesis compensate for P-eIF2α inhibitory activity. DCs do not display the features of active acute or chronic ISRs, despite PERK activation, high eIF2α phosphorylation or SubAB exposure. These characteristics allow DCs to activate and polarize T Cells in stressful physiological conditions encountered during infection or different pathologies. Moreover, our results show that actin cytoskeleton reorganization regulates eIF2α phosphorylation and protein synthesis, a biochemical cross-talk also illustrated by the impairment of migration in activated PERK-deficient DCs. Our results suggest therefore that GADD34 inactivation impacts the immune-stimulatory capacity of DCs (Clavarino et al. 2012b), but not that of PERK. We were unable to confirm that the small eIF2B activator, ISRIB and PERK inactivation inhibit cytokine expression and secretion in DCs, although this was recently described for bone marrow-derived macrophages.

**Supplementary Table 1:** List of data sets used for Immune cells in the GSEA.

| Series | doi | reference | description | class | replicate |
| --- | --- | --- | --- | --- | --- |
| GSE9810 | 10.1186/gb-2008-9-1-r17 | GSM247587 | CD8alpha cDCs | cDC1 | 1 |
| GSE9810 | 10.1186/gb-2008-9-1-r17 | GSM247588 | CD8alpha cDCs | cDC1 | 2 |
| GSE9810 | 10.1186/gb-2008-9-1-r17 | GSM247589 | CD11b cDCs | cDC2 | 1 |
| GSE9810 | 10.1186/gb-2008-9-1-r17 | GSM247590 | CD11b cDCs | cDC2 | 2 |
| GSE9810 | 10.1186/gb-2008-9-1-r17 | GSM247591 | pDCs | pDC | 1 |
| GSE9810 | 10.1186/gb-2008-9-1-r17 | GSM247592 | pDCs | pDC | 2 |
| GSE9810 | 10.1186/gb-2008-9-1-r17 | GSM247593 | NK cells | NK | 1 |
| GSE9810 | 10.1186/gb-2008-9-1-r17 | GSM247594 | NK cells | NK | 2 |
| GSE9810 | 10.1186/gb-2008-9-1-r17 | GSM247595 | B lymphocytes | B | 1 |
| GSE9810 | 10.1186/gb-2008-9-1-r17 | GSM247596 | B lymphocytes | B | 2 |
| GSE9810 | 10.1186/gb-2008-9-1-r17 | GSM247597 | B lymphocytes | B | 3 |
| GSE9810 | 10.1186/gb-2008-9-1-r17 | GSM247598 | CD8 T lymphocytes | T_CD8 | 1 |
| GSE9810 | 10.1186/gb-2008-9-1-r17 | GSM247599 | CD8 T lymphocytes | T_CD8 | 2 |
| GSE2389 | 10.1016/j.immuni.2005.01.016 | GSM44979 | CD4 T | T_CD4 | 1 |
| GSE2389 | 10.1016/j.immuni.2005.01.016 | GSM44982 | CD4 T | T_CD4 | 2 |

**Supplementary Table 2:**
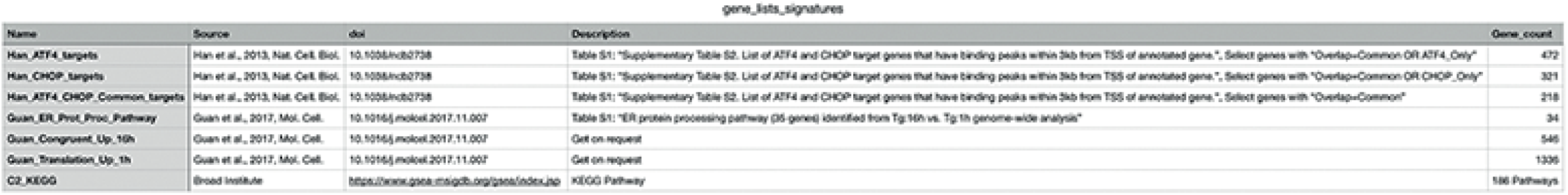
List of data sets used for ATF4 and clSR in the GSEA.

**Supplementary Table 3.**
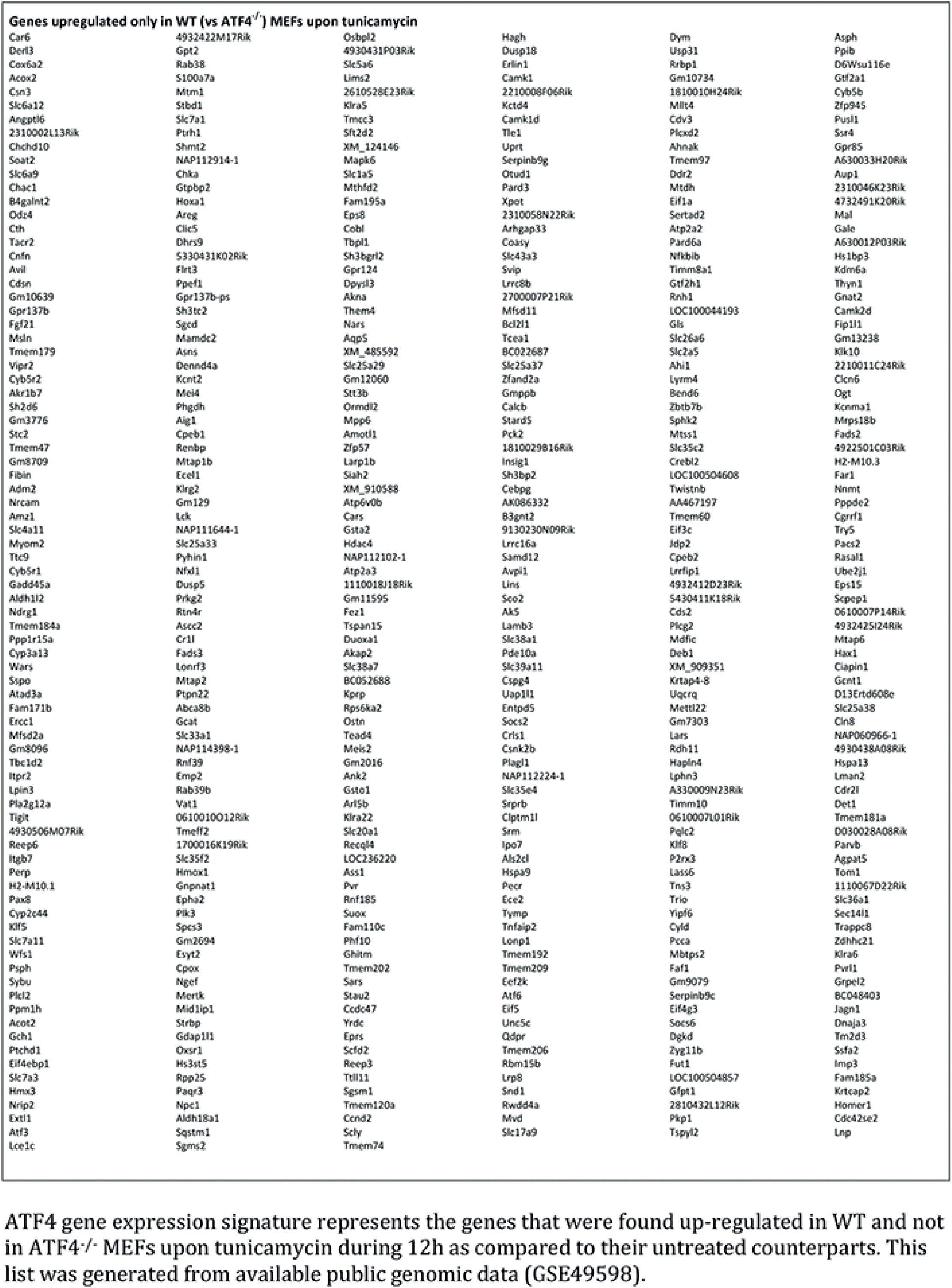

